# Uncovering Plaque-Glia Niches in Human Alzheimer’s Disease Brains Using Spatial Transcriptomics

**DOI:** 10.1101/2024.09.05.611566

**Authors:** Denis R. Avey, Bernard Ng, Ricardo A. Vialle, Nicola A. Kearns, Katia de Paiva Lopes, Artemis Iatrou, Sashini De Tissera, Himanshu Vyas, Devin M. Saunders, Daniel J. Flood, Jishu Xu, Shinya Tasaki, Chris Gaiteri, David A. Bennett, Yanling Wang

## Abstract

Amyloid-beta (Aβ) plaques and surrounding glial activation are prominent histopathological hallmarks of Alzheimer’s Disease (AD). However, it is unclear how Aβ plaques interact with surrounding glial cells in the human brain. Here, we applied spatial transcriptomics (ST) and immunohistochemistry (IHC) for Aβ, GFAP, and IBA1 to acquire data from 258,987 ST spots within 78 postmortem brain sections of 21 individuals. By coupling ST and adjacent-section IHC, we showed that low Aβ spots exhibit transcriptomic profiles indicative of greater neuronal loss than high Aβ spots, and high-glia spots present transcriptomic changes indicative of more significant inflammation and neurodegeneration. Furthermore, we observed that this ST glial response bears signatures of reported mouse gene modules of plaque-induced genes (PIG), oligodendrocyte (OLIG) response, disease-associated microglia (DAM), and disease-associated astrocytes (DAA), as well as different microglia (MG) states identified in human AD brains, indicating that multiple glial cell states arise around plaques and contribute to local immune response. We then validated the observed effects of Aβ on cell apoptosis and plaque-surrounding glia on inflammation and synaptic loss using IHC. In addition, transcriptomic changes of iPSC-derived microglia-like cells upon short-interval Aβ treatment mimic the ST glial response and mirror the reported activated MG states. Our results demonstrate an exacerbation of synaptic and neuronal loss in low-Aβ or high-glia areas, indicating that microglia response to Aβ-oligomers likely initiates glial activation in plaque-glia niches. Our study lays the groundwork for future pathology genomics studies, opening the door for investigating pathological heterogeneity and causal effects in neurodegenerative diseases.

## Introduction

The most striking pathologic features of Alzheimer’s disease (AD) are lesions known as amyloid-beta (Aβ) plaques. Activated microglia and reactive astrocytes often cluster around and within Aβ plaques and jointly influence Aβ processing and plaque biology^1^. In AD animal models, the recruitment and activation of glial cells around Aβ plaques are often associated with local inflammatory response, but whether they serve a neuroprotective, reparative, or neurotoxic function is still debated^2,3^. In postmortem human brains, high-resolution image analysis shows that microglia and astrocytes form reactive glial nets around Aβ plaques, especially around compact plaques^4^. Furthermore, the glial response near dense core plaques progresses after the Aβ load reaches a plateau^5^. Notably, microglia and astrocytes demonstrate remarkably different transcriptional signatures in human AD brains from those observed in mice^6–8^ .Thus, a critical question is how activated glial cells interact with Aβ plaques in plaque-glia niches to affect local cellular composition and gene expression in human brains.

Recent omics studies identified genes and gene networks associated with Aβ plaques at bulk and single-cell/nuclei (sc/sn) levels^6–11^. The step needed to advance the field in this space is to connect local transcriptomic differences with local tissue pathology in the spatial context. Recent advances in spatial profiling technologies enable genome-wide molecular measurements while preserving the spatial pattern of cells and pathologies. Such technologies have recently been applied in AD to reveal gene expression and cell composition changes near pathologies, but these studies primarily focus on AD mouse models^12,13^. An additional Spatial Transcriptomics (ST) study on postmortem human brains revealed altered co-expression patterns of gene modules near AD pathologies, but this study only comprised a handful of samples^14^.

Here, we apply ST and immunohistochemistry (IHC) to 78 human postmortem brain sections from 21 individuals to obtain 258,987 spatially resolved transcriptomic profiles and paired pathology measurements. With this large and unique dataset, we aim to decipher the interaction effects of Aβ plaques and reactive glia by asking two questions: 1) what the transcriptomic responses are associated with Aβ plaques in human brains, and 2) to what extent do plaque-surrounding glia modify these responses. Leveraging the intensity measurements of Aβ, GFAP, and IBA1, we stratify the ST spots into four major types of plaque-glia niches and untangle their differences in gene expression, cellular composition, and intercellular communication. We then validated the apoptotic and neuronal effects of Aβ and glial activation on postmortem brain sections using IHC. Our results show an exacerbation of synaptic and neuronal loss in low-Aβ or high-glia areas, indicating that multiple glial cell states, especially activated MG states, contribute to local immune response.

## Results

### Spatial Transcriptomics on postmortem DLPFC of 21 ROSMAP participants

We obtained 1 cm^3^ blocks of fresh-frozen tissue from the dorsolateral prefrontal cortex (DLPFC) of 21 female ROSMAP participants, 13 with AD and 8 with no cognitive impairment (NCI) (Fig.1A; Supplementary Table 1). We sectioned tissue coronally to acquire up to four 10-***μ***m sections per brain for 10X Visium ST, and for each ST section, two adjacent sections for IHC staining. For ST sections, we performed H&E staining, brightfield imaging, ST library construction, and sequencing based on the 10X protocol (Fig. 1A). In total, we obtained spatial gene expression from 258,987 spots and detected an average of 3,505 genes and 9,908 transcripts/unique molecular identifiers (UMIs) per spot (Supplementary Fig. 1A-C).

**Figure 1.**
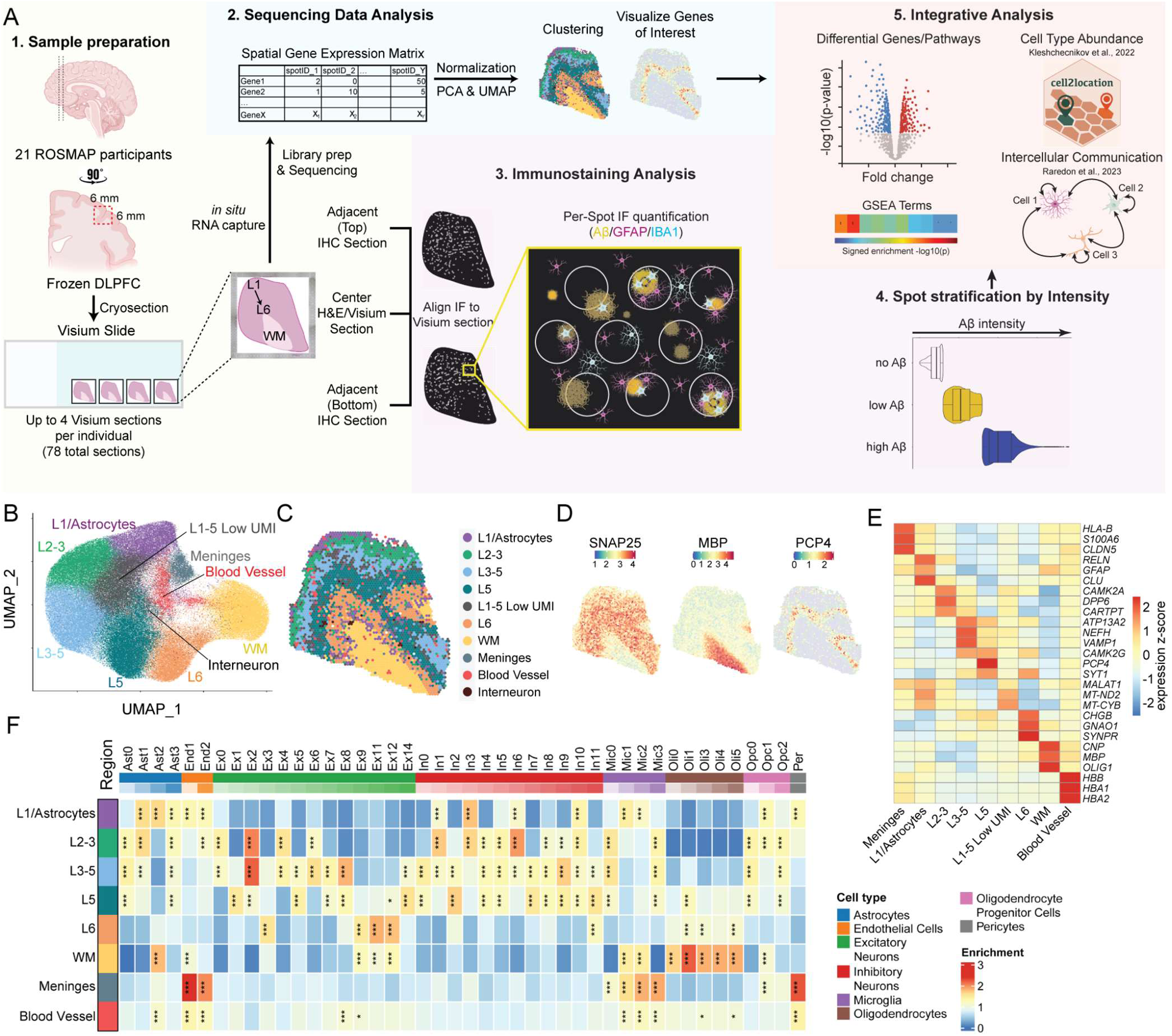
Spatial Transcriptomics Profiles of Postmortem DLPFC from 21 individuals. (A) Diagrammatic summary of experimental workflow. Per individual, up to four 10-μm frozen sections of DLPFC were subjected to spatial transcriptomic sequencing using Visium. For each ST section, two adjacent sections were stained for DAPI, Aβ, GFAP, and IBA1 and imaged by fluorescence microscopy. IHC images were then aligned to the middle ST section, enabling spot-level quantification of IHC intensity for each stain and subsequent identification of genes associated with amyloid plaques and/or glial reactivity. Cartoon graphics created with BioRender.com (B) UMAP visualization of all ST spots from all individuals. Clusters were annotated manually based on the enriched expression of known layer- and/or cell-type-specific genes. (C) Spot clusters overlaid on a representative section (Section 8_C). (D) The expression level of *SNAP25* (pan-neuronal), *MBP* (oligodendrocytes; white matter), and *PCP4* (Cortical layer 5) overlaid on a representative tissue section (Section 8_C). (E) Heatmap showing z-score of average expression of a subset of cluster-enriched genes. (F) Heatmap of GSEA normalized enrichment score (NES) for cell2location predicted neural sub-cell-types abundance across all spots comprising each spot cluster (*p < 0.01, **p < 0.001, ***p < 1e-4).

To determine the layer identities of ST spots, we first normalized the ST data using SCTransform^17^ to account for technical artifacts and applied Harmony^15^ to correct for sample-specific differences. Note that we used SCTransformed Harmony-integrated expression values only for spot clustering. We then performed dimensionality reduction and unsupervised clustering with the Louvain algorithm to identify clusters of spots with similar transcriptional profiles. We manually annotated the clusters based on known layer-specific marker genes (Fig. 1B, 1E; Supplementary Table 2). As expected, the spots clustered by cortical layer or tissue region (Fig.1C), with marker genes, such as *SNAP25*, *MBP*, and *PCP4,* highly expressed in gray matter (GM), white matter (WM), and layer 5 (L5), respectively (Fig.1D). The proportions of spots assigned to the clusters are independent of AD status and are relatively consistent across individuals (Supplementary Fig. 1F-J). The cluster-enriched genes overlap well with those found in bulk RNA-seq and other ST studies on human DLPFC^16–18^ (Supplementary Fig. 1K).

Using a published single-nuclei RNA-seq dataset of the DLPFC^9^ as a reference of 44 brain cell types, we further applied cell2location^19^ to estimate cell type abundance in each spot (Supplementary Table 3). We found that our annotated clusters comprise the expected major cell types (Fig. 1F). For example, WM spots are enriched for oligodendrocytes; meninges and blood vessel spots are enriched for endothelial cells and pericytes; and cortical layers are enriched for neuronal subtypes (Fig. 1F).

As a data quality check, we compared the co-expression patterns of our ST data against that of bulk RNASeq data from the DLPFC of 478 individuals^11^. Specifically, we generated pseudobulk estimates of gene expression by summing the UMIs across all spots of each individual and applied the Speakeasy algorithm^20^ to cluster genes into co-expression modules (Supplementary Fig. 2A). We identified 23 modules ranging from 30 to 1,199 genes per module. All ST modules are preserved in the bulk dataset and vice versa (Supplementary Fig. 2B-C). Gene set enrichment analysis (GSEA)^21^ shows that all 23 modules are enriched for relevant biological function and/or neural cell types (Supplementary Fig. 2D; Supplementary Table 4).

### ST spot stratification by Aβ and glial staining

We collected two adjacent sections for each ST section and stained them for Aβ, GFAP, and IBA1. We used GFAP and IBA1 because studies show that increased expression of GFAP^22^ and IBA1^6^ well correlate with astrocyte reactivity and microglia activation, respectively, near Aβ-amyloid plaques. After imaging the stained sections, we performed background correction using BaSIC^23^ to account for field-specific technical artifacts in establishing a uniform baseline of signal intensity. We then aligned each pair of adjacent IHC images to the middle ST section and quantified the average fluorescence intensity per spot for each antibody staining. To assess the accuracy of the alignment, we correlated GFAP gene expression from ST data and protein expression from IHC data across all spots for each section (Supplementary Fig. 3A-B).

To investigate the effects of Aβ plaques, we first stratified the 258,986 ST spots into three groups: no, low, and high Aβ, based on the average fluorescence intensity of Aβ in the corresponding 55-μm-diameter area from the two adjacent IHC stained sections (Fig. 2A-B, Methods). Specifically, spots with log_2_(average Aβ intensity + 1) values between 2.5 and 4 were classified as no Aβ, between 4 and 6.5 as low Aβ, and above 6.5 as high Aβ. These cutoffs were selected based on manual scoring of 781 plaques. A threshold of 6.5 (between low and high Aβ) maximized the distinction between diffuse and compact/dense-core plaque types. Under these cutoffs, the high Aβ spots are more enriched for compact and dense core plaques. In contrast, the low Aβ spots are more enriched for diffuse plaques (*OR*=3.02, *p*=3.21e-9, Fig. 2C). Consistently, a recent vibrational imaging study also demonstrated that increasing Aβ contents alongside the ascending plaque stages from diffuse to compact to dense core^24^. To validate this result, we performed IHC on fixed DLPFC sections from 9 AD ROSMAP participants and manually annotated 722 plaques (Supplementary Fig. 4A). Compared to fresh-frozen sections used for ST, these FFPE sections have better preservation of tissue morphology, cleaner antibody signal, and increased image resolution. We stratified the 722 plaques based on Aβ intensity into low (bottom 75%) or high (top 25%) groups (Supplementary Fig. 4B). Consistent with our observation on frozen sections, we found that low-Aβ areas are enriched for diffuse plaques, while high-Aβ areas are enriched for compact and dense core plaques (*OR*=14.98, *p*=4.89e-38, Supplementary Fig. 4C). We then further stratify the low and high Aβ into glia-high spots [defined by log_2_(average intensity + 1) of IBA1 and/or GFAP >6.8 for IBA1 and/or >7.2 for GFAP] and glia-low spots [defined by intensities for both IBA1 and GFAP< 6.2 for IBA1 and <6.5 for GFAP]. Manual assessment of glia-high/low spots indicated that glia-high spots typically contain greater than 4 glial cells (astrocytes expressing GFAP and/or microglia expressing IBA1), while glia-low spots contain 0-4 glial cells (Fig. 2D; Supplementary Fig. 5).

**Figure 2.**
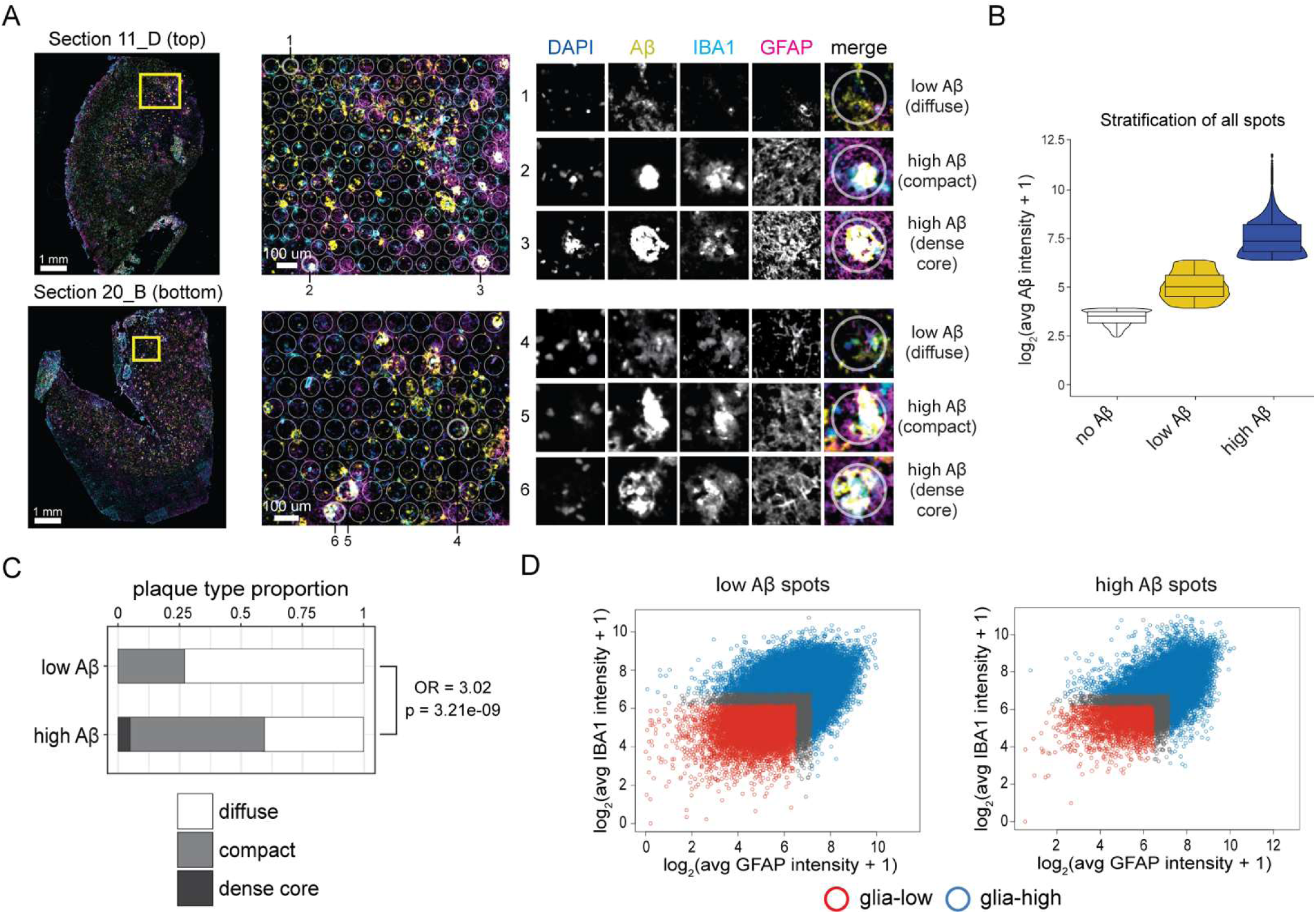
Stratification of tissue microdomains by adjacent-section IHC intensities. (A) Representative images of different plaque types from two ST-adjacent sections. Six spots (1-6; including all major plaque types) are shown with a 55-µm spot area superimposed to scale. (B) We stratified all ST spots by Aβ intensity into no (white; log_2_(avg Aβ intensity +1) between 2.5 and 4), low (yellow; log_2_(avg Aβ intensity +1) between 4 and 6.5), or high (blue; log_2_(avg Aβ intensity +1) > 6.5) groups. The distribution of Aβ IF intensity is shown for each group. (C) A subset of 781 spots from ST-adjacent sections were selected for manual plaque-type annotation (Supplementary Table 5). The proportion of plaque types is shown, stratified by low or high intensity of Aβ (OR: odds ratio). (D) Spots were further stratified by the abundance of glia (GFAP: x-axis, IBA1: y-axis). Scatterplots show the average GFAP and IBA1 intensities for each ST spot among Low Aβ (left) and High Aβ (right) groups (red: glia-low, blue: glia-high). Gray-colored spots had intermediate GFAP/IBA1 intensity and were not sorted into a group.

### Low-Aβ spots show transcriptomic profiles indicative of a more neurotoxic local environment

To examine the transcriptomic effects of Aβ, we first summed the expression values of all gray matter spots within a brain section for each stratified spot group (Fig. 3A) to generate a pseudobulk matrix for that given section. We then normalized the pseudobulk estimates across brain sections using TMM-voom^25^. To contrast the normalized pseudobulk expression of the stratified spot groups, we applied linear mixed models (LMM) with false discovery rate (FDR) correction for all tested genes and contrasts. We accounted for age, RNA Integrity Number (RIN), and batch as fixed effects and within-subject correlation as a random effect. Contrasting expression levels of low Aβ against high Aβ spots, we detected 93, 140, and 159 differentially expressed genes (DEGs) for combined, glia-low, and glia-high conditions (FDR-adjusted *p*<0.05; Fig. 3B-D; Supplementary Table 6). Many DEGs are unique for a specific contrast, with only 18 genes overlapping among all three contrasts. These results indicate that low Aβ spots display different local transcriptomes compared to high Aβ spots, and the amount of surrounding reactive glia modify local responses.

**Figure 3.**
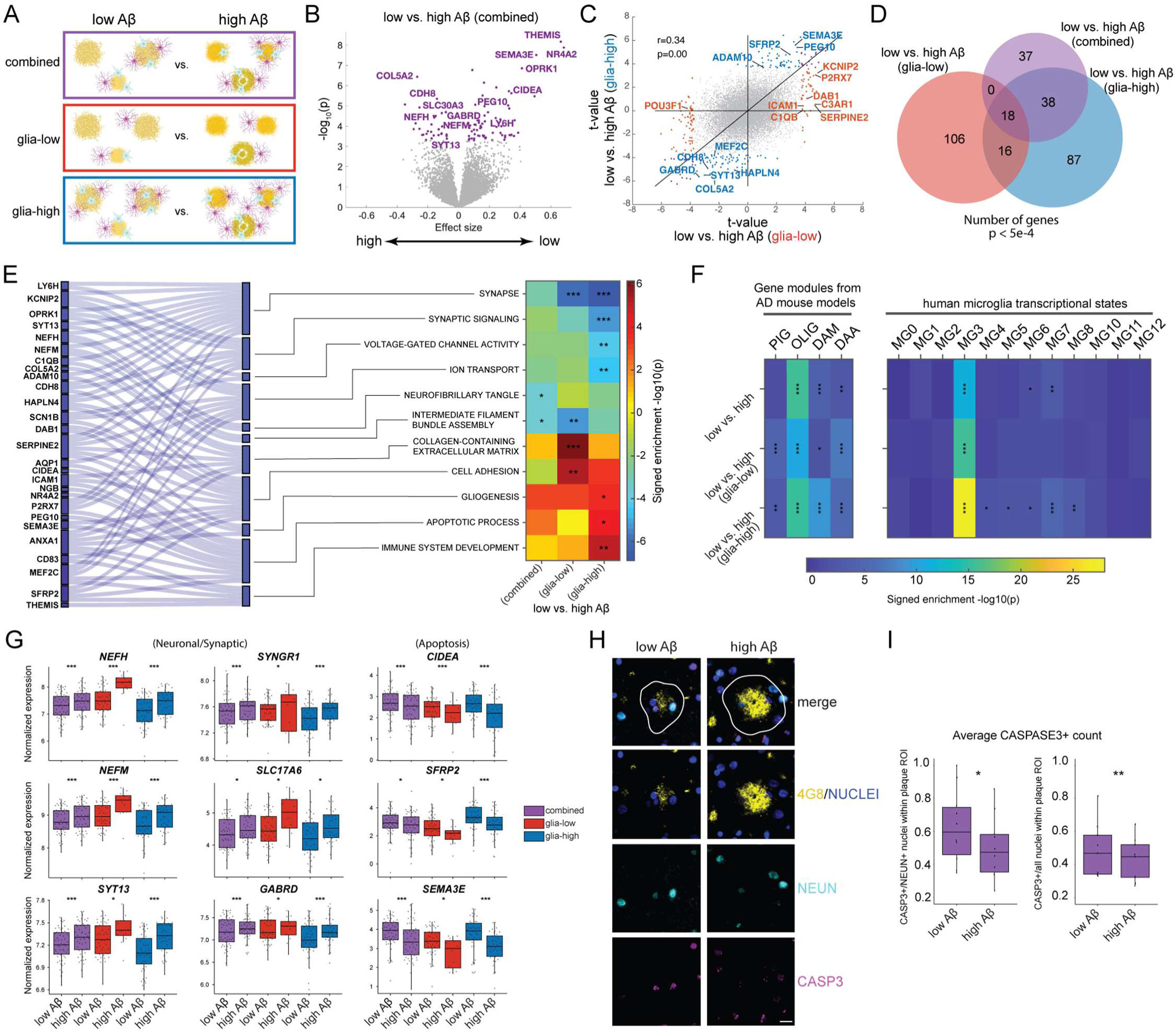
Differential effects of low vs. high Aβ on the local transcriptome. (A) Schematic of the three contrasts tested. Aβ and glia graphics created with BioRender.com (B) Volcano plot of low vs. high Aβ. Genes with FDR-adjusted p<0.05 in purple. (C) Scatterplot of Aβ effects stratified by glia-low or glia-high. Genes with FDR-adjusted p<0.05 in red and blue for glia-low and glia-high conditions, respectively. (D) Venn diagram showing the number of DEGs found across Aβ contrasts. (E) GSEA enrichment of Aβ contrasts showing GO terms and canonical pathways (*p<5e-4, **p<5e-5, ***p<5e-6) (F) GSEA enrichment of Aβ contrasts for relevant genesets (*p<5e-2, **p<5e-3, ***p<5e-4). PIG^26^: plaque-induced genes, OLIG^26^: oligodendrocyte gene module, DAM^27^: disease-associated microglia, DAA^28^: disease-associated astrocytes; MG0-12: microglia states in human AD brains^8^. (G) Boxplots of representative genes for each Aβ contrast (*p<0.05, ***p<5e-4). (H) Images of representative plaques from FFPE tissue sections stained with DAPI (blue), Aβ (4G8; yellow), NEUN (cyan), and cleaved Caspase 3 (magenta) (scale bar = 25 um). (I) Average number of Caspase 3 puncta within NEUN+ nuclei (left) or total nuclei (right) in the area within and surrounding plaques. Points represent average values from >100 ROIs for each of 10 AD individuals (*p=0.041, **p=2.61e-4).

GSEA on the low versus high Aβ contrasts shows reduced synaptic functions (Fig. 3E), especially under the glia-high condition. Notably, the downregulation of intermediate filament assembly and upregulation of the ECM/cell adhesion pathways are more pronounced under the glia-low condition. By contrast, the downregulation of ion transport and upregulation of immune and apoptosis pathways are more pronounced under the glia-high condition. In addition, we observed a negative enrichment of genes associated with neurofibrillary tangles.

Furthermore, we performed GSEA with specific genesets relevant to plaque-glia niches (Fig. 3F), including previously reported multicellular gene module of plaque-induced genes (PIG)^26^, gene module changes related to oligodendrocyte (OLIG) response^26^, disease-associated microglia (DAM)^27^, and disease-associated astrocytes (DAA)^28^ detected in mouse AD models, as well as different microglia states identified in human AD brains^8^. We detected a positive enrichment of PIG, OLIG, DAM and DAA under both glia-low and glia-high conditions, indicating some degree of preservation in response to disease pathology between humans and mice. Our result also showed a positive enrichment for OLIG signature under all glia conditions, concordant with the previous mouse ST study showing that OLIG module is highly expressed in the mild Aβ domains but decreases in microenvironment with dense Aβ pathology^26^. Among the reported human AD microglia states^8^, we detected the strongest enrichment for the MG3 signature under all glia conditions, especially under the glia-high condition. MG3 microglia were reported as “ribosome biogenesis” MG state, which most resembles mouse DAM. MG3 DAM microglia also express inflammatory genes enriched for cytokine production, antigen presentation, and microglial activation^8^. Of note, we also detected an enrichment of reported MG4 (lipid processing), MG5 (phagocytic), MG6 (stress signature), MG7 (glycolytic), and MG8 (inflammatory III) for the low Aβ glia-high condition, suggesting that microglia activation states contribute to the differential responses between low and high Aβ spots.

At the gene level, we detected lower expression of neuronal (*NEFH, NEFM*) and synaptic (*SYT13, SYNGR1, SLC17A6, GABRD*) genes, and higher expression of apoptosis (*CIDEA*) genes in the low Aβ spots across all the glia conditions (Fig 3G). The expression of extracellular matrix (ECM)- and immune-related genes (*ICAM1*, *SERPINE2*, *DAB1*) are higher in the low Aβ spots under the glia-low condition (Supplementary Fig. 6A). By contrast, the expression of apoptosis/immune-related genes (*SEMA3E*, *ADAM10*, *SFRP2*) are higher in the low Aβ spots under the glia-high condition (Fig. 3G, Supplementary Fig. 6). These results indicate that the low Aβ spots represent a more neurotoxic local environment than high Aβ spots, and plaque-associated reactive glia modulate pathogenic mechanisms. To validate the observed differences in the apoptosis process between low and high Aβ spots, we conducted IHC on FFPE brain sections from 10 additional AD individuals with antibodies against NeuN and cleaved caspase-3. For Aβ staining, we generate a region of interest (ROI) for each 4G8+ staining object. Consistent with our ST IHC stratification criteria, the top 25% of 4G8-based objects based on the scaled average 4G8 intensity was defined as high Aβ and the lower 75% as low Aβ. We collected ∼517 ROIs for cleaved Caspase-3 from those 10 AD individuals. The ROI was defined as the nuclei+/NeuN+ areas within each 4G8 object and a 25 μm ring around it. To contrast the effects of low vs. high Aβ on apoptosis, we applied a negative binomial mixed model (NEBULA) with Caspase-3 count as the response, NeuN+ nuclei count as the offset, age at death, and sex as fixed effect confounding factors and modeled within-subject correlations among ROIs from the same subject with a random effect. We also tested all nuclei count (DAPI+) as the offset. We detected more cleaved caspase-3 puncta co-localized with NeuN+ nuclei and total nuclei near low Aβ plaques compared to high Aβ plaques (Fig. 3H-I).

### Low Aβ spots show cell composition and intercellular communication indicative of neurodegeneration

To examine differential Aβ effects on cell composition, we quantified the abundance of 44 brain cell types in each spot using cell2location^19^ and contrasted no Aβ, low Aβ, and high Aβ spots (Fig. 4A). To calculate representations of each cell type, we aggregated and normalized the predicted number of cells by section and spot group. We then constructed LMM to test for differential cell abundances between stratified spot groups (Supplementary Table 7). In general, we observed more cell-type composition differences between low versus no Aβ than high versus low Aβ, indicating a more profound impact on cell-type proportions during initial Aβ accumulation in aged brains. Specifically, we observed a negative enrichment of most inhibitory neuron (In) subtypes (In1, In3, In5, In6, In8, In10) and all oligodendrocyte progenitor cell (Opc) subtypes (Opc0, Opc1, Opc2) (FDR adjusted *p*<0.05; Fig. 4A-C) for low vs no Aβ, in line with literature that loss of inhibition and aberrant OPC disruption are early pathological signs in AD^29–31^. In addition, we observed a negative enrichment of End2, Ex0, Ex4, Mic2, Per in low versus no Aβ condition and a negative enrichment of Ex11, Ex12, In2, In4, Olig0, Olig1, Olig3, and Olig5 in the high versus low Aβ condition (Fig. 4A-C). These results indicate a selective cell vulnerability across all major cell types. The early depletion of OPCs may contribute to the reduction of oligodendrocytes at the later stage. Notably, we did not detect cell abundance differences for most astrocytes and microglia, and one possible explanation is that proliferation and cell death may co-occur for the glia surrounding Aβ plaques, rendering a relatively constant cell representation. The positive enrichment of most post-mitotic neurons likely results from the selective loss of vulnerable cells, leaving those relatively more resilient neurons enriched.

**Figure 4.**
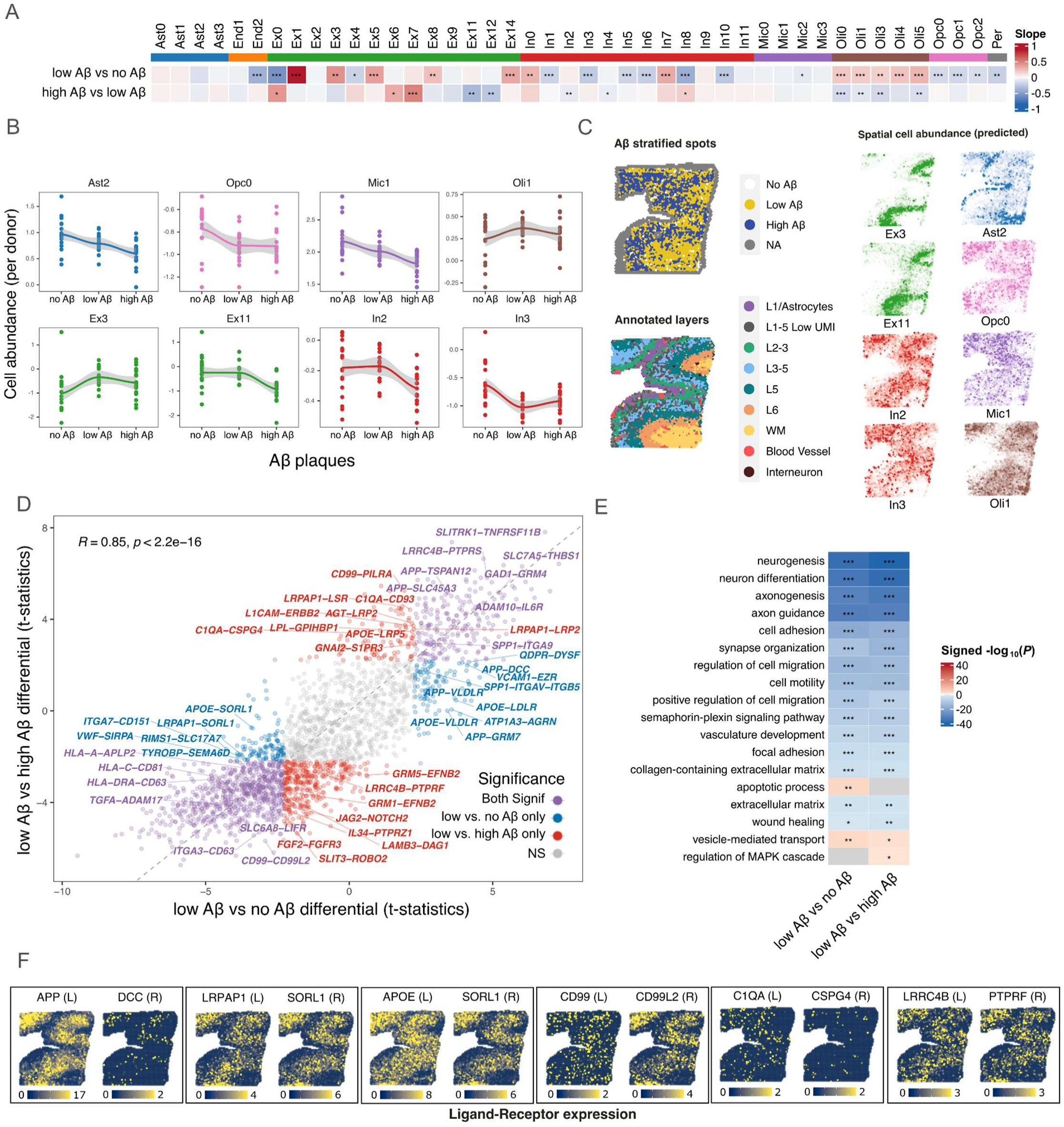
Aβ plaques influence cell composition and intercellular communication. (A) Cell abundance differences in response to Aꞵ. The heatmap shows slopes of cell abundance differences between no Aꞵ vs. low Aꞵ and low Aꞵ vs. high Aꞵ. The number of cells predicted for each spot was aggregated by section (median) and quantile-normalized. A linear mixed model was applied, comparing normalized abundances in each group while controlling by donor (as a random effect) and age. *P*-values are adjusted by FDR (*p ≤ 0.05, **p ≤ 0.01, ***p ≤ 0.001). (B) Cell dynamics in response to Aꞵ. Line plots showing selected sub-cell type abundances by stratified Aꞵ spots. The y-axis represents the quantile-normalized number of cells aggregated by section (median), and the x-axis represents spots grouped by Aꞵ stratification. (C) Example of the section colored by Aꞵ stratification spots, layer annotation, and number of predicted cells for selected sub-cell types. (D) Differential ligand-receptor (LR) interaction genes between low Aꞵ vs. high Aꞵ (y-axis) and low Aꞵ vs. no Aꞵ spots (x-axis). Each dot corresponds to an LR pair tested; the axis represents the t-statistics from linear model tests accounting for repeated donors. Significant LR pairs (FDR < 0.05) are highlighted in red (low Aꞵ vs. high Aꞵ only), blue (low Aꞵ vs. no Aꞵ only), or purple (significant in both comparisons). (E) Selected Gene Ontology terms enrichment from LR differentially expressed in low Aꞵ vs. no Aꞵ and low Aꞵ vs. high Aꞵ. Both ligand and receptor genes were considered for the enrichment analysis. P-values were adjusted *g:SCS* method from *gprofiler*. Colors in the heatmap represent the signed -log10(adj. P). *p ≤ 0.05, **p ≤ 0.01, ***p ≤ 0.001. (F) Representative section (Section 15_D) image colored by normalized expression of selected differentially expressed LR gene pairs. All the statistical analyses were performed on gray matter spots only.

We next investigated differential Aβ effects on intercellular communications using NICHES^32^, an algorithm that allows the embedding of ligand–receptor (LR) signal proxies to estimate the local microenvironment for each spot in ST data. We first conducted low versus no Aβ and low versus high Aβ differential analysis to identify differentially expressed LRs (Supplementary Table 8). We noticed a positive association (R = 0.85) between these contrasts. Many LR pairs were dysregulated in both contrasts, with functions related to synapses, neuron differentiation, and ECM pathways (Fig. 4D-F). These results indicate that the low Aβ spots constitute a more neurotoxic microenvironment than high Aβ spots. Consistently, low Aβ spots presented genes related to greater apoptotic processes compared with no Aβ spots. In addition, low Aβ spots also show LRs (*APP-DCC*, *APP-VLDLR*, *APOE-LDLR*) related to greater vesicle-mediated transport compared with no Aβ and high Aβ spots (Fig. 4D-F), suggesting a potential compensatory mechanism for Aβ clearance. In line with our cell abundance result, our analysis suggests that low Aβ spots present more profound LR differences related to neuronal and synaptic degeneration than high Aβ spots, highlighting the importance of early intervention during plaque accumulation. In addition, low Aβ spots were also enriched for LRs related to greater Mitogen-Activated Protein Kinase (MAPK) cascades and immune pathways (*C1QA-CD93, CD99-PILRA*) compared with high Aβ spots. Immune modulation has been an active topic for AD therapies. On the other hand, Aβ has been shown to activate the MAPK, and inhibiting MAPKs reduces Aβ generation and alleviates cognitive impairments in AD models^33^. Our results corroborate those studies, suggesting that combining immune modulation and MAPK inhibition may provide an effective therapeutic approach to alleviate Aβ neuronal damage, especially in early AD.

### Plaques with surrounding glia display transcriptomic differences indicative of elevated neuronal toxicity

To test whether plaque-surrounding glia influences the local transcriptome within plaque niches, we contrasted the glia-high versus glia-low gray matter spots with and without stratification by Aβ intensity (Fig. 5A). We first estimated TMM-normalized pseudobulk expression of glia-high and glia-low spots for each condition and then applied LMM with FDR correction for all tested genes and contrasts. We detected 230, 241, and 63 DEGs for combined, low Aβ and high Aβ conditions (FDR adjusted *p*<0.05; Fig. 5B-D; Supplementary Table 9), respectively. As expected, we detected higher expression of glial markers (*SPARC*, *CD44*, and *HLA-DRA*) in the glia-high spots across all three Aβ conditions (Fig. 5B,G). Furthermore, we detected lower expression of excitatory and inhibitory neuronal makers (*RORB*, and *PVALB*), synaptic signaling (*SYT2, SLC6A17, GRM7, and GABBR2*), ion channels (*SCN1B*), and neurofilaments (*NEFH/M/L*) in glia-high spots (Fig. 5B, G). We also detected higher expression of translation genes (*EEF1A1*) in glia-high spots (Fig. 5B, Supplementary Fig. 7), consistent with a recent study demonstrating that increased local protein synthesis in microglia serves as a necessary step for phagocytosis^34^. Our GSEA analysis showed a negative enrichment for synapses, exocytic vesicles, neural filaments, and voltage-gated channel activity and positive enrichment for many immune-related terms under all Aβ conditions (Fig. 5E), in line with the recent high-resolution imaging study demonstrating that reactive glia nets around Aβ plaques are toxic inflammatory microenvironments in postmortem AD brains and animal models^4^.

**Figure 5.**
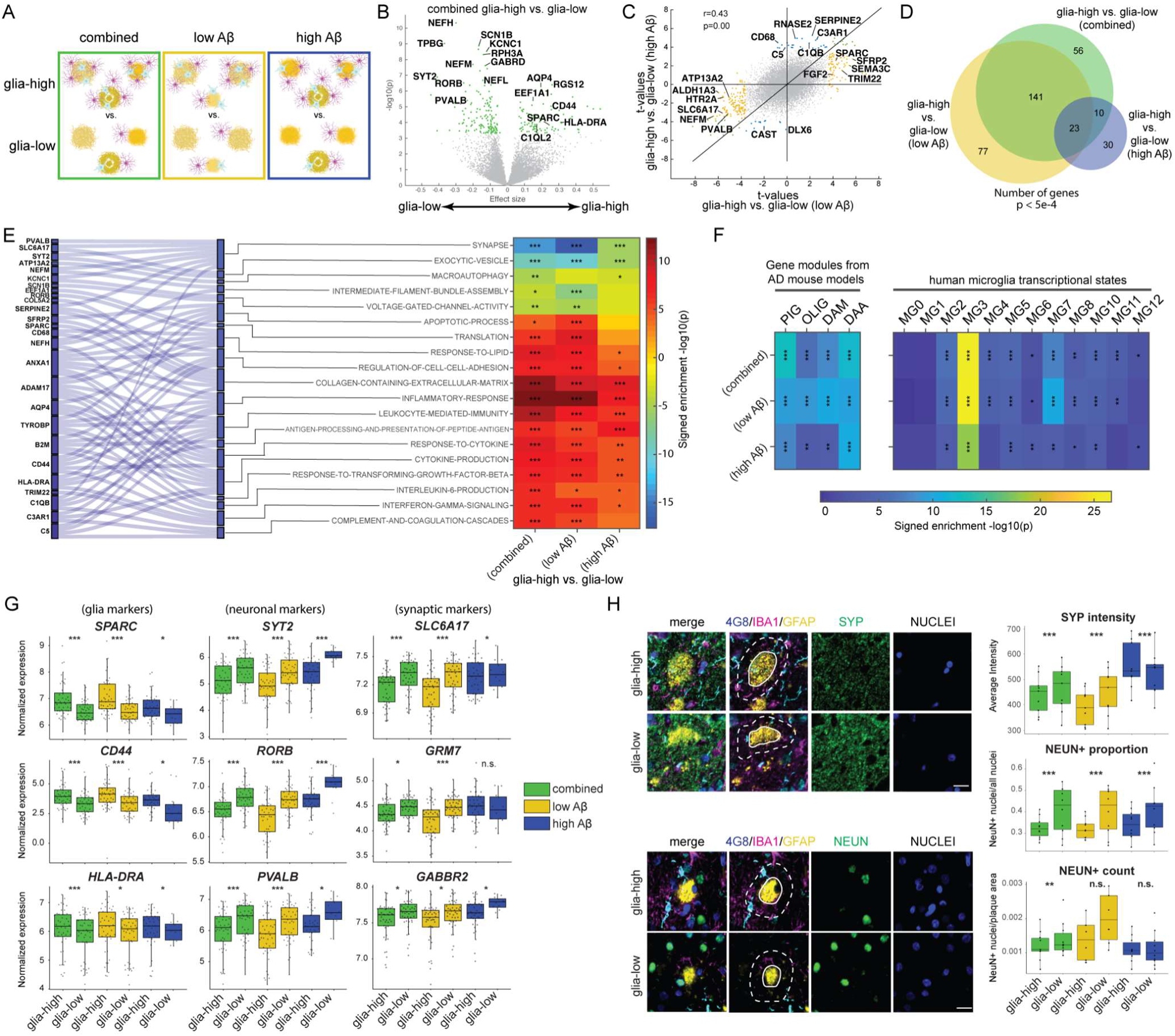
Differential effects of glia abundance on the local transcriptome. (A) Schematic of the three contrasts tested. Aβ and glia graphics created with BioRender.com (B) Volcano plot of glia-high vs. glia-low. Genes with FDR-adjusted p<0.05 in green. (C) Scatterplot of glia effects stratified by low or high Aβ. Genes with FDR-adjusted p<0.05 in yellow and blue for low and high Aβ conditions, respectively. (D) Venn diagram on the number of DEGs found across glia contrasts. (E) GSEA enrichment of glia contrasts for GO terms and canonical pathways (*p<5e-4, **p<5e-5, ***p<5e-6). (F) GSEA enrichment of glia contrasts for Aβ/glia-relevant genesets (*p<5e-2, **p<5e-3, ***p<5e-4). PIG^26^: plaque-induced genes, OLIG^26^: oligodendrocyte gene module, DAM^27^: disease-associated microglia, DAA^28^: disease-associated astrocytes; MG0-12: microglia states in human AD brains^8^. (G) Boxplot of representative genes found for each glia contrast (*p<0.05, ***p<5e-4, n.s. not significant). (H) Images of representative plaques from FFPE tissue sections stained with DAPI (blue), Aβ (4G8; yellow), IBA1 (cyan), GFAP (magenta), and synaptophysin (SYP; green, top-left panel) or NEUN (green, bottom-left panel). SYP and NEUN display lower-intensity staining surrounding glia-high plaques compared to glia-low plaques (quantification on right). Points represent average values from each of 10 AD individuals quantified in the area surrounding plaques (scale bar = 25µm; **p<0.005, ***p<1e-10, n.s. not significant).

Furthermore, we detected specific gene changes and geneset enrichment for the different Aβ conditions. Under the low Aβ condition, expression of genes related to lysosome/autophagy (*ATP13A2*) is lower, and expression of genes related to immune (*TRIM22*) and axon guidance/apoptosis (*SEMA3C*) is higher in glia-high spots (Fig. 5E, Supplementary Fig. 7). Under the high Aβ condition, expression of genes related to phagocytosis (*CD68*), ECM (*SERPINE2*), and complement pathway (*C5*) are higher in the glia-high spots (Fig. 5E, Supplementary Fig. 7A). Our GSEA analysis showed a stronger negative enrichment of synapses, intermediate filament assembly, and voltage channels in the low Aβ condition, accompanied by stronger positive enrichment of apoptosis, complement cascades, and cytokine production, supporting a more profound effect of surrounding glia in the low Aβ condition. Thus, our results indicate that plaque-surrounding glia may elevate neuronal toxicity and synapse degeneration in the aged human brain, especially in low-Aβ plaque niches. Because inflammatory response^35^, complement cascades^36^, disturbed extracellular vesicle release^37^, and abnormal lysosome-autophagy pathways^38^ have been implicated in AD pathogenesis, our results also suggest that the plaque-surrounding glia are implicated in multiple facets of AD neurodegeneration.

In addition to gene ontology (GO), we further applied GSEA with gene signatures of PIG^26^, OLIG^26^, DAM^27^, and DAA^28^ and different microglia states^8^ (Fig. 5F). We detected enrichment of PIG, OLIG, DAM, and DAA under both low Aβ and high Aβ conditions, indicating an elevated glial response from multiple glial cell types, including microglia, astrocytes, and oligodendrocytes. Indeed, the PIG module has been shown to involve multiple cell types and represent intercellular crosstalk between astrocytes and microglia^12^. Among the reported human AD microglia states^8^, we detected enrichment for almost all MG states except for MG0 (homeostatic) and MG1 (neuronal surveillance)^8^. The MG3 DAM (ribosome biogenesis) state showed the strongest enrichment under all Aβ conditions. The enrichment of the MG3 state for glia response is consistent with the positive GO enrichment for translation (Fig. 5E). In addition, the enrichment for MG4 (lipid processing) and MG7 (glycolytic) aligned with the positive GO enrichment for lipid response. Further, the enrichment for MG5 (phagocytic), MG8 (inflammatory II), and MG10 (inflammatory III) are in line with the GO enrichment for immune responses. As such, these results indicate that diverse activated MG states, especially MG3, are present around the plaques, contributing to local immune response. Taken together, our results suggest that multiple glial cell states arise around plaques and contribute to local immune response.

We then set out to validate the synaptic and neuronal loss in the high-glia spots. To this end, we stained FFPE sections from 10 AD individuals with Aβ, GFAP, and IBA1 and synaptophysin (*SYP*) to label synapses or NeuN to label neuronal nuclei (Fig. 5H). Consistent with our ST IHC analysis, 4G8 objects with an ROI scoring in the top 25% for scaled GFAP or IBA1 average intensities were defined as glia-high and the remaining objects as glia-low. The intensity and count measurements were collected from the ROI defined as the 25 μm ring around the plaque. We then applied the NEBULA model with NeuN count as the response to test the effects of glia-high vs. glia-low on neuron loss. We also examined glia effects on synapse loss by applying linear mixed models with average SYP as the response and within-subject correlations among ROIs from the same subject as a random effect. We detected reduced staining of these neuronal/synaptic markers near glia-high plaques compared with glia-low, especially for low Aβ plaques (Fig. 5H), in line with our ST data (Fig. 5G). Notably, our result is consistent with a recent ST study showing alterations in GABAergic and glutamatergic signaling corresponding to microglia density in the plaque niche in a mouse model of AD^39^. In addition, we stained the brain sections with antibodies against Aβ, GFAP, IBA1, and CD68 to examine whether glia-high Aβ spots are associated with increased neuroinflammation. Our result shows a higher abundance of CD68 protein near glia-high than glia-low plaques (Supplementary Fig. 7C).

### Plaques with surrounding glia affect cell composition and intercellular communication

To examine whether plaque-surrounding glia alter the cell composition around plaques, we contrasted cell abundance for each cell type between glia-low and glia-high groups for low Aβ, high Aβ, and combined conditions in gray matter (Supplementary Table 7). As expected, we observed a positive enrichment of most astrocytes, microglia, and OPCs subtypes in glia-high spots across all three Aβ conditions (Fig. 6A-C). Glia-high spots are also accompanied by similar negative enrichment of neuronal subtypes, including Ex1, Ex5, Ex7, In0, and In7 (Fig. 6A-C). Notably, these neuronal subtypes are distinct from those identified from Aβ contrasts, indicating that neuron subtypes may differ in their vulnerability to Aβ toxicity and inflammatory response depending on the presence of surrounding glia. We also detected a positive enrichment of some types of post-mitotic neurons, likely resulting from the selective loss of vulnerable neurons (Fig. 6A-C).

**Figure 6.**
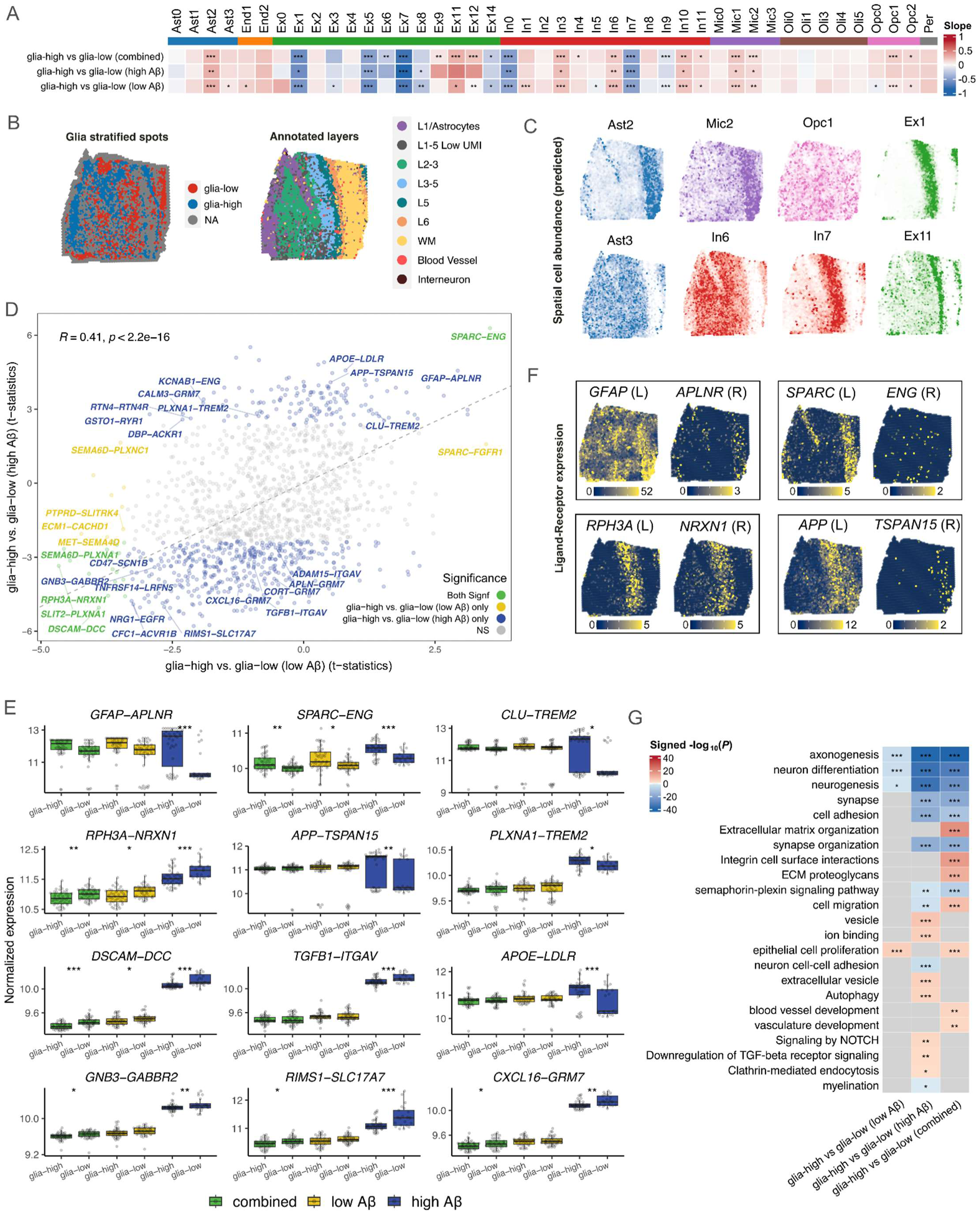
Plaque-surrounding reactive glia modifies cell composition and intercellular communication. (A) Cell abundance differences in response to glia. The heatmap shows slopes of cell abundance differences between glia-high vs. glia-low under low Aꞵ, high Aꞵ, or combined. The number of cells predicted for each spot was aggregated by section (median) and quantile-normalized. A linear mixed model was applied, comparing normalized abundances in each group while controlling by donor (as a random effect) and age. *P*-values are adjusted by FDR (*p < 0.05, **p< 0.01, ***p < 0.001). (B) Representative section (Section 15_D) colored by glia stratification spots and layer annotation. (C) Representative section colored by the number of predicted cells for selected sub-cell types. (D) Differential ligand-receptor (LR) genes between glia-high vs. glia-low under High Aꞵ spots (y-axis) and glia-high vs. glia-low under low Aꞵ spots (x-axis). Each dot corresponds to an LR pair tested; the axis represents the t-statistics from linear model tests accounting for repeated donors. Significant LR pairs (FDR < 0.05) are highlighted in blue (glia-high vs. glia-low under low Aꞵ only), yellow (glia-high vs. glia-low under high Aꞵ only), or green (significant in both comparisons). (E) Normalized expression levels by stratified spot groups for selected differentially expressed LR pairs. (*p ≤ 0.05, **p ≤ 0.01, ***p ≤ 0.001). (F) Example sections colored by normalized expression of selected differentially expressed LR gene pairs. All the statistical analyses were performed on gray matter spots only. (G) Selected Gene Ontology terms enrichment from LR differentially expressed between glia-high vs. glia-low under low Aꞵ, high Aꞵ, and combined were considered for the enrichment of both ligand and receptor genes. P-values were adjusted using the *g:SCS* method from *gprofiler*. Colors in the heatmap represent the signed -log10(adj. *P*) (*p ≤ 0.05, **p ≤ 0.01, ***p ≤ 0.001).

We then applied NICHES to estimate differences in microenvironments centering around each ST spot. We first contrasted glia-high versus glia-low to identify differentially expressed LRs under low Aβ, high Aβ, and combined conditions (Supplementary Table 9). We observed a positive correlation (R = 0.41) between the low Aβ and high Aβ conditions. The expression of LRs related to synaptic pathways (*RPH3A-NRXN1*, *DSCAM-DCC, GNB3-GABBR2, RIMS1-SLC17A7, and CXCR16-GRM7*) was downregulated in the glia-high spots across all Aβ conditions (Fig. 6F-G). In the high Aβ condition, we also detected an upregulation of genes related to immune (*CLU-TREM2*, *PLXNA1-TREM2*, *APOE4-LDLR* and *GFAP-APLNR*) and extracellular cell matrix (*SPARC-ENG*), vesicle transport (*APP-TSPAN15*) as well as a downregulation of TGF-β1 signaling (*TGFB1-ITGAV*). Concordantly, our GSEA analysis on differentially expressed LRs revealed more substantial LR differences in the high Aβ condition, including increased autophagy as well as reduced cell adhesion and myelination (Fig. 6D-G). Because NICHES measure LR interactions of the microenvironment surrounding each ST spot, the more pronounced functional implications in the high Aβ condition may reflect a culmination of progressive plaque development, glial activation, and neurodegeneration.

### ST glial response to Aβ plaques recapitulated by co-cultured microglia-like cells upon Aβ oligomer treatment

Studies have shown that Aβ plaques are potential reservoirs of oligomeric Aβ, often forming a “halo” around Aβ plaques^40,41^. To examine whether the glial transcriptional signature detected by ST reflects response to Aβ oligomers and to what extent microglia or/and astrocyte mediate this response, we established a co-culture system encompassing iPSC-derived microglia-like cells, astrocytes, and neurons to mimic the multicellular nature of the brain tissue (Fig. 7A). We performed immunofluorescent staining 72h after co-culture and detected IBA1+ microglia-like cells (iMGLs), GFAP+ astrocytes, and DCX+ neurons (Fig. 7B). We then treated co-cultures with Aβ oligomers for 24 hours and harvested the cells for scRNA-seq using the 10X Genomics Chromium platform.

**Figure 7.**
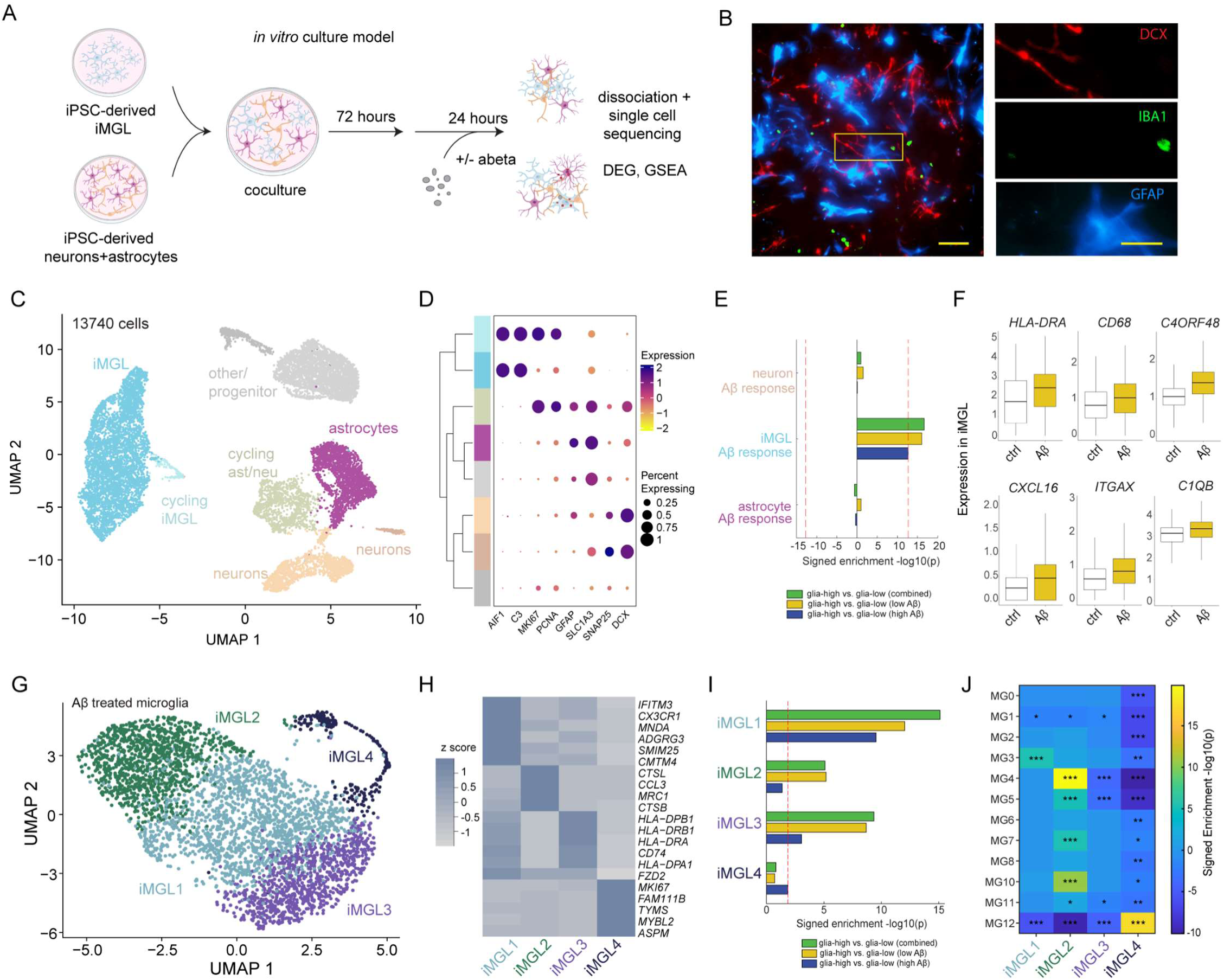
Aβ oligomer treatment of co-cultured iMGL partially recapitulates ST glia response to Aβ plaques. (A) Schematic of *in vitro* co-culture experiments. Graphics created with BioRender.com (B) Representative immunofluorescence for DCX (neuronal), IBA1 (iMGL) and GFAP (astrocyte) proteins in co-culture models, scale bar: 100µm, inset: 50µm. (C) UMAP visualization of cells from control and Aβ oligomer-treated cultures, colored by annotated cell type clusters. (D) Dotplot of cluster/cell-type enriched genes. (E) GSEA enrichment of Aβ-induced genes for ST glial response by cell type. Red dotted line corresponds to FDR-threshold. (F) Boxplots of genes differentially expressed in iMGL upon Aβ treatment. (G) UMAP visualization of Aβ oligomer-treated iMGL, colored by subtype cluster. (H) Heatmap of the top 5 differentially expressed gene markers per iMGL subtype cluster. (I) GSEA enrichment of Aβ-treated iMGL cluster DEGs (Supplementary Table 11) for ST glial response. Red dotted line corresponds to FDR-threshold. (J) GSEA enrichment of Aβ-treated iMGL cluster (vs. control) DEGs (Supplementary Table 12) for reported MG state genesets in human AD brains^8^ (*p<5e-2, **p<5e-3, ***p<5e-4).

We acquired scRNAseq data from 13,740 cells under the Aβ untreated and treated conditions. After standard data processing and filtering, we applied unsupervised clustering on the first 15 principal components to group the cells (Supplementary Fig. 8C). We identified 5 major cell clusters that we annotated based on classical markers as iMGLs, astrocytes, neurons, cycling neuroglial progenitors, and other progenitors (Fig. 7C-D). To examine the Aβ oligomers’ effect, we analyzed differential gene expression for neurons, astrocytes, and iMGLs, and identified 199, 58, and 271 Aβ significant DEGs, respectively (Supplementary Table 10). We then performed GSEA on the glia-high versus glia-low contrasts from ST data with the DEGs upregulated upon Aβ-oligomer treatment as the geneset. We detected a positive enrichment of the ST glial response in iMGLs. However, no enrichment was observed in astrocytes and neurons (Fig. 7E). The DEGs in iMGLs included many immune, complement cascade, and phagocytosis genes, such as *HLA-DRA*, *CD68*, *C4ORF48*, *CXCL16*, *ITGAX*, and *C1QB* (Fig. 7F). In agreement, GSEA on the co-culture data revealed enrichment for increased immune response, reduced cell metabolism, and cytokine release, aligning with the functional pathways of the ST glial response (Supplementary Table 10). Thus, our results suggest that the molecular effect of plaque-surrounding glia in tissue may result from the Aβ oligomers residing in plaques. Although both astrocytes and microglia are present around plaques, our results support the idea that microglia may play a primary role in mediating glial response. Consistently, the recent spatial profiling on plaque niches in mouse AD brains and high-resolution imaging on the human brain reveal that microglia closely surround plaques and astrocytes are more distant from the plaques^4,42^.

We then further divided the Aβ-treated iMGLs into 4 subclusters (iMGL1-4; Fig. 7G, 7H) and applied GSEA on the glia-high vs glia-low contrasts from ST data with DEG markers of each iMGL subcluster as genesets (Supplementary Table 11). iMGL1, iMGL2, and iMGL3 showed different degrees of enrichment of *the ST* glial response (Fig. 7I). Specifically, iMGL1 demonstrated enrichment under both Aβ-high and Aβ-low conditions, whereas iMGL2 and iMGL3 showed enrichment primarily in the Aβ-low condition (Fig. 7I). These results suggest that the responses of iMGL1-3 to oligomeric Aβ, especially iMGL1, constitute a major element of the ST glial response. To understand which MG states the iMGL subclusters resemble in human AD brains, we applied GSEA on the transcriptomic signature of each iMGL subcluster with markers of 12 reported MG states as genesets. (Supplementary Table 12)^8^. Notably, iMGL1 is positively enriched for the MG3 DAM state, which is the most enriched state in the ST glial response (Fig. 5F). iMGL2 is enriched for MG4 (lipid processing), MG5 (phagocytic), MG7 (glycolytic), and MG10 (inflammatory III) (Fig. 7J), which is also in line with the enriched states in the ST glial response (Fig. 5F). Thus, our results show that the MG states in the AD brain can be recapitulated by co-cultured microglia upon Aβ treatment. This suggests multiple reported MG states, especially MG3, shape the local glial response to Aβ deposition in human AD brains.

## Discussion

While Aβ plaques and reactive glia have long been known to be associated with neurodegeneration, their interaction effects on local transcriptomes have never been studied at scale using ST. Here, we report 258,986 microdomains with paired ST-IHC measurements across 78 samples from 21 ROSMAP individuals. This large sample size enables statistical inference of Aβ and glial effects that are not merely observed in a limited sample size. By coupling ST readouts with IHC features of Aβ plaques and glial activation, we identify genes, pathways, cell type composition, and cell interaction differences associated with distinct plaque-glia niches, shedding light on the cellular and molecular environments of plaque-related neurodegeneration. We demonstrate that low-Aβ areas are more neurotoxic than high-Aβ areas and an increased abundance of reactive glia in plaque-glia niches worsens neurodegeneration. The glial response genes we identified by ST are enriched for multiple MG states identified by snRNA-seq of human AD brains. Importantly, exposure of co-cultured iMLCs to Aβ oligomers recapitulates the ST glial transcriptional response, as well as the reported MG states in human AD brains.

### Low versus high Aβ spots

By contrasting low Aβ and high Aβ spots, we detected transcriptomic changes indicative of increased synaptic degeneration in the low Aβ spots. First, our DEG and GSEA analysis identified genes and functional pathways related to synaptic degeneration. Second, our cell2location^19^ analysis infers more profound cell representation differences in the low Aβ plaque spots. Third, our NICHES analysis reveals more drastic LR differences related to neuronal and synaptic degeneration in the low Aβ spots. Fourth, our IHC validation experiment shows increased cleaved caspase-3 puncta co-localized with NeuN+ nuclei and total nuclei near low Aβ plaques compared to high Aβ plaques. Thus, all our results point toward more detrimental effects in low Aβ spots. Our result resonates with a recent study showing that Aβ can accumulate into plaque depositions inside damaged neurons, subsequently contributing to extracellular Aβ plaque formation^43^, suggesting that the neuronal damage observed in AD could occur at the early stage of plaque development when Aβ loads are relatively low. This observation is also concordant with a recent label-free vibrational imaging study showing that early-stage plaques consist mainly of toxic Aβ oligomers and protofibrils, whereas late-stage plaques primarily contain Aβ fibrils^24^. Moreover, a recent electron cryomicroscopy (cryo-EM) study demonstrated two types of Aβ42 filaments in human brains. Type I filaments are mostly in sporadic AD cases with abundant compact plaques, and Type II filaments are in familial AD cases with abundant diffuse plaques^44^. Considering the general severity of Aβ deposition in familial AD, this result also supports more neuronal toxicity of diffuse plaques.

Another interesting observation gleaned from contrasting low Aβ and high Aβ spots is that surrounding glial cells influence the local transcriptome. For example, we detected an upregulation of ECM pathways in the glia-low condition and an upregulation of immune and apoptotic pathways in the glia-high condition (Fig. 3E, 3G). Since low-Aβ plaques represent the early phase of plaque development, this divergence could represent differential coping mechanisms in the glia-high versus glia-low spots. In the glia-low condition, the upregulation of *ICAM* and *DAB1* (Supplementary Fig. 6A) might be beneficial because *ICAM1* can protect neurons against Aβ by blocking NF-κB signaling^45^, and *DAB1* activation by Reelin (*RELN*) is known to modulate synaptic activity^46,47^. In addition, the upregulation of *RELN and DAB1* has also been associated with vulnerable neurons of AD brains^48^, indicating that the protective effects of Reelin and DAB1 might be eventually overwhelmed by disease progression. In the glia-high condition, the upregulation of *ADAM10* and *SFRP2* could also be compensatory. *ADAM10* in astrocytes has been shown to cleave the APP protein and prevent Aβ generation^49^, and secreted frizzled-related proteins (SFRPs) can bind to Aβ and prevent protofibril formation, thereby modulating astrocyte-microglia crosstalk in neuroinflammation^50–52^. These results indicate that the plaque niches might utilize different mechanisms to cope with the Aβ-related cell damage depending on the context of glial activation.

### The effects of plaque-surrounding glia on the local niche

To examine the effect of surrounding glia in the local plaque niches, we contrasted glia-high versus glia-low spots in the low Aβ and high Aβ conditions. Our analysis indicates that the surrounding glia may mount a robust inflammatory response and exacerbate neurodegeneration, especially in the low Aβ spots. Our IHC validation experiments further validated this result, showing reduced neuronal/synaptic and increased inflammation marker expressions near glia-high plaques compared with glia-low plaques (Fig. 5H). The neuronal degeneration in glia-high spots is consistent with recent studies. First, microglia contain and accumulate Aβ aggregates, which could lead to microglial death and the release of intracellular Aβ aggregates, contributing to plaque expansion and neurodegeneration^53^. Indeed, eliminating microglia in AD mouse models stops plaque growth and reverses AD-related transcriptomic changes^54^. Second, microglia activate the NLRP3 inflammasome, which subsequently drives the formation and release of ASC specks (apoptosis-associated speck-like protein containing a CARD), increasing Aβ oligomer formation, immune response, and neuronal damage^55^. Lastly, dysregulated crosstalk between microglia and astrocytes also escalates neuroinflammation and cell death. For example, A1 astrocytes can be induced by activated microglia, which can induce neuronal death and synaptic loss; in accordance, the presence of A1 astrocytes is reported in human AD brain^56^. Thus, our results support a harmful and non-resolving inflammatory response in glia-surrounding plaques, resulting in elevated neuronal degeneration.

Furthermore, our GSEA analysis shows that the ST glial response is enriched for signatures of reported mouse gene modules of plaque-induced genes (PIG), oligodendrocyte (OLIG) response, disease-associated microglia (DAM), and disease-associated astrocytes (DAA), as well as different microglia (MG) states identified in human AD brains, indicating that multiple glial cell states arise around plaques and contribute to local immune response. Among all the enriched MG states, MG3 is the most enriched in the ST glial response. This MG state has been shown to resemble mouse DAM and expresses inflammatory genes enriched for cytokine production, antigen presentation, and microglial activation^8^. The strong enrichment of the MG3 state provides a potential therapeutic cell state target for fine-tuning the immune response in human AD brains. In addition, our GSEA analysis also shows a greater leukocyte-mediated immunity in glia-high spots across all Aβ conditions (Fig. 5E), which is unexpected as the innate immune response (MGs and astrocytes) has been considered the primary culprit of neuroinflammation in AD. Nevertheless, our result aligns with recent studies demonstrating that microglia can mediate T cell infiltration in the mouse brain with tauopathy, and the number of T cells was correlated with the extent of neuronal loss. An increased number of T cells was also found in the AD postmortem brains^57^. In addition, CD163-positive monocyte-derived cells were shown to infiltrate brain parenchyma near Aβ plaques, contributing to plaque-associated myeloid cell heterogeneity^58^. Therefore, an immune hub involving innate and adaptive responses may be present in aged human AD brains, especially in the plaque-glia niches.

### *In vitro* validation of ST glia response

The iPSC-derived microglia have been shown to recapitulate disease microglia states upon exposure to AD-related stimuli and master regulators^8,59^. To disentangle the glia effect observed in brains, we established an iPSC-derived multicellular culture system that emulates the cellular interactions in the brain tissue microenvironment. We show that iMCLs, upon Aβ-oligomer treatment, but not astrocytes, mirror the ST glial response. Among iMGL subclusters, iMGL1-3 clusters demonstrate an enrichment for glial response. Notably, iMGL1 most resembles MG3 DAM state, and iMGL2 resembles multiple MG states in human AD brains. These results further extend our ST analysis of the glial response, indicating that multiple MG states mediate glial response and microglia’s response to toxic Aβ oligomers initiates glial activation in the plaque-glia niches. Although we did not detect enrichment for other cell types, the crosstalk among astrocytes, neurons, and microglia may contribute to microglia’s strong molecular response. Indeed, crosstalk between astrocytes and microglia has been shown to increase the degradation of Aβ aggregates in iPSC-derived cultures^60^. Furthermore, given that we only treated co-culture for 24h, it is also possible that prolonged treatment may trigger a pronounced response in neurons and astrocytes directly or indirectly. Nonetheless, the positive enrichment of the ST glial response in iMGLs supports a primary effect of microglia in driving glial response in plaque-glia niches.

### Limitations

There were several limitations of this study. First, our ST samples were collected from aged female individuals (mean age of 92). Therefore, whether our results can generalize to younger age groups or males remains to be determined. Second, because Aβ spots primarily came from AD cases, we were underpowered to assess the intersection of AD status with the detected transcriptional signatures in plaque-glia niches. Third, as the Visium platform is at ∼55-μm resolution and both astrocytes and microglia accompanied most plaque-containing spots, we were also underpowered to analyze astrocyte- and microglia-specific transcriptional responses in plaque niches. Fourth, due to frequent freezing artifacts of postmortem frozen tissue and the large size of this dataset (258,986 spots), we could not definitively categorize some of the plaques as the diffuse, compact, or dense core; thus, we used no, low and high Aβ to group the ST spots instead. Finally, because of the relatively low resolution of the Visium platform and the limitation to choosing three proteins for IHC, we focused on extracellular Aβ pathology and surrounding glia to investigate the Aβ-glia niches. Future studies will investigate molecular responses to intracellular pathologies, such as neurofibrillary tangles, using high-resolution ST platforms.

### Conclusions and Future Directions

Our study demonstrates how advanced ST technology can be combined with classical histology to answer questions that neither alone can address. Our ST workflow and computational framework represent a new and accessible approach to investigating pathology heterogeneity in AD and other neurodegenerative diseases. By coupling ST information with traditional morphological features of plaques, we provide novel insights into brains’ cellular and spatial context, linking the abundance of plaques and nearby reactive glia to neuron vulnerability, synapse degeneration, and inflammatory response. Our study also demonstrates the power of integrating ST, snRNA-seq, and human cell models to decipher the sequential events and cell states in plaque-glia niches, indicating that diverse MG activation states arise around Aβ plaques and microglial response to toxic Aβ oligomers initiates glial activation. Overall, our study lays the groundwork for future pathology genomics, opening a new avenue for addressing the pathology heterogeneity in neurodegenerative diseases. Future studies with increased sample size and diversity, high-resolution spatial profiling platforms, the ability to work with fixed/archived tissues, and co-detection of protein(s) on the same section will ameliorate many technical issues we encountered and enable finer delineation of cell-type-specific responses to distinct pathologies.

## Acknowledgments

We are grateful to those who agreed to donate their brains for research. We thank all the employees at RADC for their support and assistance. This study was supported by NIA grants R01AG074082 and R01AG079223 (to Y.W.), P30AG10161, P30AG72975, R01AG015819, R01AG017917, and U01AG61356 (to D. A. B.), and R01AG061798 (to C.G.).

## Author contributions

D.R.A., S.D.T., and D.J.F. performed sample prep and ST experiments. J.X. performed ST data and scRNA-seq data processing. B.N, R.V., D.R.A, N.K., A.I., K.L., and S.T. performed data analysis. N.K. and D.S. performed cell culture experiments. H.V. and D.S. performed immunohistochemistry experiments. D.R.A., B.N., R.V., and Y.W. wrote the manuscript with input from all co-authors. D.A.B runs the parent ROSMAP study. Y.W. conceived and supervised this study. All authors reviewed and approved the final manuscript.

## Lead Contact

Further information and requests may be directed to the corresponding author Yanling Wang

## Materials Availability

This study did not generate new unique reagents.

## Data Availability

Sample/spot information, differentially expressed genes, differentially expressed ligand-receptors, gene modules, ST, and scRNA-seq cluster marker genes are provided in the Supplementary Information. Raw and normalized count matrices of the Spatial Transcriptomics and ST-paired IHC image data are available at Synapse under accession code syn53141470.

## Online Methods

### Tissue collection for Spatial Transcriptomics

The human brain tissues were obtained through the ROSMAP (Religious Orders Study and Memory and Aging Project) cohort studies at the Rush Alzheimer’s Disease Center, Rush University Medical Center, Chicago, IL. Written informed consent was given by the donors for brain autopsy and for the use of material and clinical data for research purposes, in compliance with national ethical guidelines. Clinicopathological information of the donor, including postmortem time, age, and sex, is recorded (Supplementary Table 1). Neuropathological diagnosis of the Aβ deposition, neurofibrillary tangles, and other pathologies is provided (Supplementary Table 1). We selected 13 individuals clinically diagnosed as AD with high pathology and 8 controls with no cognitive impairment (NCI) and no or very little pathology. All individuals are female and matched for age. We chose all female individuals because there were insufficient males in the ROSMAP cohort matching our selection criteria (e.g. RIN > 5, age-matched, minimal freezing artifacts in frozen DLPFC). Frozen brain blocks of the right-hemisphere DLPFC from 21 individuals were cut (∼ 1 cm^3^), transferred on dry ice, and stored at -80C.

Frozen brain blocks were embedded in OCT and cryosectioned coronally to a thickness of 10 μm using a Leica CM1950 cryostat set at -17C. Sectioning was done in sets of three (middle section for ST and two adjacent sections for IF), with 10-50 μm between each set. Sections were attached to Visium slides and stored at -80C in a slide mailer for up to one week before proceeding with spatial transcriptomics. ST-adjacent sections were attached to pre-labeled Leica superfrost plus slides and stored at -80C in slide mailers for up to two weeks before proceeding with immunohistochemistry.

### Spatial Transcriptomics

Spatial Transcriptomics was performed according to the Visium Spatial Gene Expression User Guide (10X Genomics; CG000239). Briefly, cryosectioned tissue on Visium slides was transferred from -80C to the lab on dry ice and quickly thawed for 1 min on a heat block at 37C, then immediately transferred to methanol pre-equilibrated to -20C and fixed at -20C for 30 min. Sections were then stained with hematoxylin and eosin, and brightfield images were acquired with a Nikon Ti2 with NIS-Elements AR 5.11.01 64-bit software. After imaging, tissues were immediately permeabilized in 0.1% pepsin diluted in 0.1 M HCl for 18 min at 37C. Reverse transcription, second-strand synthesis, cDNA amplification (15 cycles), and library construction were performed according to the manufacturer’s instructions. For the first four individuals (#2, 8, 12, and 18), protocol version ’Rev A’ was followed, and the sample index PCR was 12 cycles. For the remaining samples, protocol version ’Rev D’ was followed, and the sample index PCR was 13-14 cycles. Library concentration was quantified by Qubit 1x dsDNA HS using a SpectraMax M3 plate reader. Library concentration and size distribution were assessed by Fragment Analyzer (Agilent). Sequencing was performed in three batches on an Illumina NovaSeq, targeting a minimum of 50,000 raw reads per spot.

### Immunofluorescence assay of ST-adjacent sections

We first attempted to perform IHC staining on the ST section, following 10X Genomics recommendations. However, due to the modified protocol required to preserve RNA integrity (i.e., methanol instead of PFA fixation, no antigen retrieval, shorter blocking, and antibody incubations), we were unable to obtain adequate staining quality. We decided instead to perform IHC on two ST-adjacent sections. Cryosectioned tissue on Leica super-frost slides was transferred from -80C to the lab on dry ice and quickly thawed for 1 min on a heat block at 37C, then immediately transferred to a slide-mailer filled with PFA pre-equilibrated to 4C and fixed at 4C for 20 min. Slides were washed twice with DPBS before performing antigen retrieval for 10 min @ 95C in 10 mM sodium citrate buffer, pH 6.0. Following antigen retrieval, slides were washed three times with DPBS, a hydrophobic barrier was drawn surrounding the tissue (two sections per slide), and then sections were blocked for 1 h at RT in blocking buffer (DPBS containing 5% normal goat serum, 1% BSA, 0.1% Tween-20, and Human TruStain FcX (Fc blocker)). Sections were then incubated overnight at 4C with primary antibodies to detect Aβ (4G8), GFAP, and IBA1 (diluted in blocking buffer). The next day, the primary antibody solution was removed, and sections were washed three times with DPBS (5 min per wash). Sections were then incubated for 1.5 h at RT in the dark with fluorophore-conjugated secondary antibodies (Alexa Fluor 488 for Aβ, Alexa Fluor 790 for GFAP, and Alexa Fluor 594 for IBA1) diluted 1:500 in blocking buffer. The secondary antibody solution was removed, and sections were washed four times with DPBS (5 min per wash), then incubated with DAPI to stain nuclei and Trueblack (according to the manufacturer’s instructions) to quench tissue autofluorescence. Slides were coverslipped and imaged on the ThermoFisher CellInsight CX7 LZR High Content Analysis Platform (HCA; 100 16-bit images per section).

### Immunostaining quantification for ST-adjacent sections

H&E images were acquired from ST sections with an Eclipse Ti2-E microscope (Nikon) at 20x magnification in 9×9 tile tiff files with 10% overlap and were stitched into a single image per section with Adobe Photoshop (23.4.2 release). Fluorescent Aβ, GFAP, IBA1, and DAPI images were acquired from the two adjacent sections with the ThermoFisher HCA at 20x magnification in 10×10 tile 16-bit tiff files. Background and shading correction were performed with the ImageJ plugin BaSiC^23^. The tiles were then stitched with a 1% overlap with the stitching plugin in ImageJ. The adjacent IHC sections were aligned to the middle/sequenced H&E section by setting 7-12 common landmarks between each adjacent image and the H&E one, then using the ImageJ plugin ’Landmark Correspondences’.

Due to the non-uniform intensity background across tiles and bright areas arising from artifacts such as tissue wrinkles and tears, we applied standard image processing techniques for further background correction and artifact removal. Using a structural element with a size well above that of amyloid plaques, microglia, and astrocytes, for each IF image, we performed grayscale opening and closing to extract large bright artifacts and local background inhomogeneity, which we regressed out from the image. With the artifact-corrected IHC images, we quantified the amount of amyloid, microglia, and astrocytes by averaging the intensity of pixels within each spot of the ST sections. We excluded spots near tissue borders overlapping with bright artifacts from our analysis.

### Immunostaining of FFPE tissue

FFPE cortical sections were pre-heated for 60 min at 60C in a dry oven and then rehydrated using a series of xylene and ethanol dilutions before rehydrating in DI water. Sections were then used for performing antigen retrieval for 20 min in boiling 10 mM sodium citrate buffer, pH 6.0. Following antigen retrieval, sections were washed with DPBS, incubated in 0.02N HCl for 20 min for peroxide blocking, washed with DPBS, and blocked for 1 h at RT in TNB blocking buffer (0.1M Tris-HCl pH 7.5, 0.15M NaCl and 0.5% Akoya blocking reagent; prepared as per manufacturer’s instructions). Sections were then incubated overnight at 4C with primary antibodies diluted in the TNB blocking buffer to detect combinations of Aβ (4G8; 1:1000), GFAP (1:1500), IBA1 (1:200), cleaved Caspase-3 (1:250), NeuN (1:100), SYP (1:500) and/or CD68 (1:100). The following day, primary antibody solution was removed from sections, washed three times with TNT wash buffer (0.1M Tris-HCl pH 7.5, 0.15M NaCl and 0.05% Tween-20) (5 min per wash) and incubated for 1 h at RT in the dark with species-appropriate fluorophore-conjugated secondary antibodies (Alexa Fluor 568, Alexa Fluor 647, Alexa Fluor 488 or Alexa Fluor 790) diluted 1:1000 in DPBS. For cleaved Caspase-3 staining, the antibody was detected with a Mouse IgM-HRP secondary, incubated for 1 hour at RT, followed by a 5 min incubation with tyramide Cy5. After secondary antibody incubation, sections were washed three times with TNT wash buffer (5 min per wash), then incubated with Trueblack for 1 min to quench tissue autofluorescence. Slides were washed two times with DPBS (5 min per wash), incubated with Hoechst 34580 (1:2000 in DPBS) for staining nuclei,mounted using Prolong antifade mountant and coverslipped. Slides were dried overnight at RT and imaged on the ThermoFisher HCA (200-300 16-bit images per section).

### Immunostaining quantification for FFPE tissue sections

IHC stained slides were imaged on the ThermoFisher HCA at 20x. 24 fields per section were selected to profile the thickness of the cortex from the meninges to the white matter for each subject. Images were processed using the compartmental analysis module in the HCA software. After background correction, 4G8 staining was used to identify plaque-like objects with the following parameters: objectarea > 20μm, smoothing = 2. The 4G8-based objects were used as a mask to generate a region of interest (ROI) for the GFAP and IBA1 channels that encompassed both the object area and an extension from the edge of each object up to 25μm without overlap. Calculated metrics, including the average ROI intensity for each channel, object size, and object shape, and counts were exported from the software. The data was filtered for an average 4G8 intensity >2000. To adjust for technical differences in staining intensities between slides, the intensity data was scaled and centered for each case, and these resulting z-scores were used for subsequent plots. To classify plaques, an investigator blinded to the HCA measurements manually scored all 4G8-based objects as diffuse, compact, or dense core plaques or as a staining artifact/vessel yielding a final sample size of n = 722 plaques from n=9 cases. For target intensity measurements, either the compartmental or colocalization analysis modules of the CellInsight system were used to define secondary objects and extract the average intensity and counts within each ROI. The top 25% of 4G8-based objects based on the scaled average 4G8 intensity was defined as high Aβ and the lower 75% as low Aβ. Objects with an ROI scoring in the top 25% for either scaled GFAP or scaled IBA1 average intensities were defined as glia-high and the remaining objects as glia-low. For NeuN (n=10)/SYP(n=9) (Fig. 5H), and CD68 (n=10) (Fig. S7C) data, intensity and count measurements were collected from the ROI defined as the 25 μm ring around the plaque (not including the plaque area). For cleaved Caspase-3 (n=10) data (Fig. 3H-I), the ROI was defined as either the nuclei+ or NeuN+ areas within each 4G8 object and a 25 μm ring around it. The average number of 4G8 based objects analyzed per case were 933 for SYP, 507 for NeuN, 1136 for CD68, and 517 for Caspase-3. To contrast the effects of low vs. high Aβ on apoptosis, we applied a negative binomial mixed model (NEBULA^61^) with Caspase-3 count as the response, NeuN+ nuclei count as the offset, age at death and sex as fixed effect confounding factors and modeled within-subject correlations among ROIs from the same subject with a random effect. We also tested all nuclei count (DAPI+) as the offset. To test the effects of glia-high vs. glia-low on neuron loss, we further applied this model with NeuN count as the response. Both nuclei count and ROI area (excluding plaques) were separately tested as the offset. Finally, we examined glia effects on synapse loss and inflammation by applying linear mixed models with average SYP and CD68 intensity, respectively, as the response, age at death and sex as fixed effect confounding factors and modeled within-subject correlations among ROIs from the same subject with a random effect.

### Spatial transcriptomics sequencing data processing

Sequencing data were pre-processed with Space Ranger pipeline (10x Genomics), which mapped the barcodes against the human genome (Gencode 27), generated FASTQ files, and generated a count matrix for each section. The individual files were then merged in a Seurat object (Seurat v4.2.0) and were processed in R. We calculated per-spot quality metrics and spots with low-measured genes (< 500) were removed. The Seurat::SCTransform function was used to normalize and scale the UMI counts based on regularized negative binomial regression. Principal component analysis (PCA) was performed on the top 3000 variable genes. Using the top 50 PCs (Supplementary Fig. 1D) and accounting for technical and biological covariates such as age, RNA integrity number (RIN), library batch, and donor, we integrated the datasets using the Harmony package (v0.1.0). We explored various tools for data integration and chose Harmony for its high performance, usability, and scalability. We only used Harmony integration to identify clusters corresponding to brain regions. All downstream analyses (differential gene expression, cell proportion estimates, and ligand-receptor analysis) were performed on pre-integration count data. After visualizing the standard deviation attributed to each Harmony embedding as an elbow plot (Supplementary Fig. 1E), we decided to use the top 10 Harmony embeddings. We applied Seurat::runUMAP to compute UMAP (uniform manifold approximation and projection) dimension reduction and Seurat::findNeighbors and Seurat::findClusters to identify clusters based on a shared nearest neighbor (SNN) clustering algorithm (Louvain clustering with resolution set to 0.3; all other parameters default). This resulted in 11 clusters, which were annotated based on their canonical markers and spatialLIBD (v7.5) to the closest anatomical cortical layer (L1, L2-3, L3-5, L5, L6), white matter (WM), meninges, and specific cellular subpopulations, namely cells with high expression of hemoglobin genes (’Blood Vessel’) and SST/NPY (’Interneuron’). The last identified cluster, which carried a glial signature, had very few spots and so was not included in downstream analyses.

### Co-expression network analysis in spatial transcriptomics

We performed co-expression network analysis using all spots and sections of the ST data. We created a pseudo-bulk gene expression matrix by summing the counts by donor. The lowly expressed genes were removed, and we kept genes with at least 1 CPM (counts per million) in all samples. Different normalization methods were tested, and TMM-voom (edgeR_3.36.0)^30^ was applied for the final matrix. To understand the major sources of variation in the gene expression matrix, we used the variancePartition^31^ R function (v1.24.0) to run a linear mixed model and attribute the percentage of variation in expression based on technical and biological variables. To minimize the effect of unwanted confounders, removeBatchEffect (Limma library v3.50.1) R function was used to adjust the expression matrix by librarybatch, RIN, age, and PMI. A total of 14,534 expressed genes from 21 donors were used as input for the SpeakEasy (SE v1.0)^21^ consensus clustering algorithm. We built the network using the results from 100 initializations of SE and Spearman correlation, which resulted in 29 modules, 23 with at least 30 genes (meaning that 99.3% of the genes were assigned to a module). The module assignment contains a list of genes by module in Supplementary Table 4.

For module functional enrichment analysis, we applied two approaches. 1) gprofiler (v2_0.2.1)^32^ with the g:GOSt R function: the webserver includes databases as Gene ontology, KEGG pathways from KEGG Reactome and WikiPathways; miRNA targets from miRTarBase and regulatory motif matches from TRANSFAC; tissue specificity from Human Protein Atlas; protein complexes from CORUM and human disease phenotypes from Human Phenotype Ontology. For multiple test corrections, we used the default g:SCS method. 2) Fisher exact test with Bonferroni correction (Adj p < 0.05) for specific gene lists as described in the Results section. All 23 modules were significantly enriched with at least one function.

For reproducibility, we ran the modulePreservation^33^ function from the WGCNA R package (v1.70-3) with our ST networks and previously published bulk RNASeq network with 478 donors. The Zsummary statistic was accessed, and a Kruskal Wallis test was applied for significance. Modules with Z < 2 were considered not preserved, between 2 and 10 preserved, and Z > 10 highly preserved (Supplementary Fig. 2B-C).

### Spot stratifications

ST spots were classified as no, low, or high Aβ based on the average fluorescence intensity of Aβ in the corresponding 55-μm-diameter area from the two adjacent immunostained sections. Spots with log_2_(average Aβ intensity + 1) less than 2.5 were not included for downstream analysis, as these were primarily regions not covered by tissue. Spots with log_2_(average Aβ intensity + 1) values between 2.5 and 4 were classified as no Aβ, between 4 and 6.5 as low Aβ, and above 6.5 as high Aβ. These cutoffs were selected based on manual scoring of 781 plaques (Fig. 2D-E). A threshold of 6.5 (between low and high Aβ) maximized the distinction between diffuse and compact/dense-core plaque types. Glia-high spots were defined by high log_2_(average intensity + 1) of IBA1 and/or GFAP (>6.8 for IBA1 and/or >7.2 for GFAP), while glia-low spots had low intensities for both IBA1 and GFAP (<6.2 for IBA1 and <6.5 for GFAP). Manual assessment of glia-high/low spots indicated that glia-high spots typically contain >4 glial cells (astrocytes expressing GFAP and/or microglia expressing IBA1), while glia-low spots contain ∼0-4 (Supplementary Fig. 5). Different glia intensity thresholds were tested, but increasing the stringency of thresholds or isolating single-positive spots (e.g. GFAP+/IBA1- or GFAP-/IBA1+) reduced the number of sections available to generate pseudobulk expression profiles, and thus reduced the power for DEG detection.

### Statistical analysis to identify Aβ and glia differential genes

For each brain section of a subject, we summed the UMI counts of all spots belonging to a plaque niche type to generate a pseudobulk estimate for each gene. We only kept expressed genes with counts per million (CPM) > 1 in at least 80% of the brain sections. With the pseudobulk estimates, we estimated the library sizes of the brain sections using the trimmed mean of M-values (TMM) of edgeR^62^ (edgeR_3.36.0) and transformed the pseudobulk estimates into log_2_ CPM using voom-limma (limma_3.50.1). For each plaque niche type, we excluded brain sections with <50 spots. To contrast the normalized pseudobulk expression of a given pair of plaque niche types, we applied linear mixed models (LMM) with plaque-glia niche types coded as a binary variable. We accounted for age, RIN, batch, and AD status as fixed effects and modeled within-subject correlation with a random effect. We declared significance at an *α* of 0.05 with false discovery rate (FDR) correction across all genes and all Aβ and glia contrasts.

For interpretation, we applied GSEA^21^ with t-values of the niche type contrasts as scores and GO terms as genesets. We also used markers of PIG^26^, OLIG^26^, DAM^27^, and DAA^28^ from mouse studies as well as different microglia states identified in human postmortem AD brain^8^ as genesets. Moreover, with our co-cultured scRNA data, we extracted marker genes or Aβ induced genes for iMGL, astrocytes, neurons and iMGL1-4 subtypes, which were further used as genesets.

### Cell abundance estimates

The abundance of specific brain cell types was estimated for each ST spot using cell2location^16^. The average expression of 44 distinct brain cell subpopulations from 8 major cell types was estimated from DLPFC snRNA-seq data^3^ and used as a reference for cell deconvolution with cell2location. We estimated the enrichment of each cell subpopulation in all annotated brain regions using GSEA. Spots in each brain region were ranked using the cell2location predicted abundance, and overrepresentation normalized enrichment scores were calculated for each cell subpopulation. *P*-values were estimated using 100 permutations and adjusted for multiple comparisons using the Benjamini-Hochberg false discovery rate (BH-FDR) correction. Relative cell abundance comparisons between stratified spots were performed by aggregating spot-level predicted cell counts into pseudo-bulk matrices by section. For each spot group, the median cell count from each of the 44 subpopulations was calculated. Only data from gray matter spots and groups with at least ten spots were included. Quantile normalization was then applied to the pseudo-bulk cell abundance matrix. Linear mixed models were used to test for differential relative cell abundances between stratified spot groups. All models were adjusted by age and repeated donor information (modeled as random effects). *P*-values were adjusted for multiple comparisons using BH-FDR correction.

### Ligand-Receptor analysis

Local microenvironment interactions were estimated from ST data using NICHES (Niche Interactions and Communication Heterogeneity in Extracellular Signaling v1.0.0)^34^. We used the Omnipath database (v3.2.8) as the ligand-receptor (LR) reference and used default parameters for data imputation, k-nearest neighbor graph (k=4), and Niche Matrix Construction (summing signaling input from Euclidean neighbors). Each section was analyzed separately, and results were merged for downstream analysis. For stratified spot differential expression comparisons, spot-level LR matrices were first averaged by section and stratification. Only data from gray matter spots and groupings with at least ten spots were included. LR genes with CPM <= 1 in < 50% of sections were removed. Then, the aggregated mean values were normalized and transformed by quantile-voom using the *limma* R package (v3.50.3). A mixed linear model was applied to compare the normalized values between each stratification group while accounting for sections derived from the same donors using the *duplicateCorrelation* function (limma_3.50.3). *P*-values were adjusted for multiple comparisons using BH-FDR correction.

### iPSC cultures and differentiation

Two iPSC lines (AICS-0090-391 purchased from the Coriell Institute and BR28^63^ obtained from Dr. Tracy Young-Pearse) were used for the culture experiments. iPSCs were maintained on Matrigel-coated dishes in mTeSR Plus (StemCell Technologies). iPSCs were passaged every 4 days with ReLeSR (StemCell Technologies) according to the manufacturer’s protocol.

To generate iMGL, we followed previously published protocols^64,65^ with minor modifications using AICS-0090-391 iPSCs. iPSC colonies were seeded at a density of 20-40 per well of a Matrigel-coated 12-well plate in mTeSR Plus supplemented with 10μM Y-27632 and cultured for 48 hours. At d0, colonies were fed with hematopoietic stem cell (iHPC) basal media [IMDM (50%), F12 (50%), insulin (0.02 mg/ml), ITSG-X (2% v/v), L-ascorbic acid 2-Phosphate magnesium (64 μg/ml), monothioglycerol (400 μM), PVA (10 ng/ml), Glutamax (1X), chemically defined lipid concentrate (1X), and non-essential amino acids (NEAA; 1X)] supplemented with FGF2 (50 ng/ml), BMP4 (50 ng/ml), Activin-A (12.5 ng/ml), and LiCl (2 mM). At d2, cultures were fed with iHPC basal media supplemented with FGF2 (50 ng/ml) and VEGF (50 ng/ml). At d4, cultures were fed with iHPC basal media supplemented with FGF2 (50 ng/ml), VEGF (50 ng/ml), TPO (50 ng/ml), SCF (10 ng/ml), IL-6 (50 ng/ml), and IL-3 (10 ng/ml) and was replaced every other day until d12. At d12, iHPCs were gently resuspended in MGL differentiation media [DMEM/F12, HEPES, insulin (0.2 mg/ml), ITS-G (2% v/v), B27 (2% v/v), N2 (0.5%, v/v), monothioglycerol (400 μM), Glutamax (1X), NEAA (1X), and additional insulin (5 μg/ml)] supplemented with M-CSF (25 ng/ml), IL-34 (100 ng/ml), and TGFβ-1 (50 ng/ml) and seeded at 2 × 10^5^ cells per well of a Matrigel-coated 6-well plate. Additional MGL differentiation media was added to cultures every two days. For the remainder of the differentiation, cells were passaged as needed to maintain a density of 2-5 × 10^4^ cells per cm^2^, retaining 50% of the media when replating. From d30-d38, MGL differentiation media was supplemented with M-CSF, IL-34, TGFβ-1, CD200 (100 ng/mL), and CX3CL1 (100 ng/mL).

To generate astrocyte/neuron cultures, we first differentiated BR28 iPSCs into neural progenitors (NPCs). iPSCs were dissociated with accutase and replated to Matrigel-coated plates at 2.5 × 10^5^ cells/cm^2^. From the following day, cultures were fed daily with neural induction media [DMEM/F12, 1X N2, 1X B27, 2mM Glutamax, 100μM NEAA, 110μM BME, 0.05% BSA] supplemented with 5μM SB432542 and 50μM LDN193189 until day 12. We then used Stem Cell Technologies astrocyte differentiation and maturation media and protocol to generate mixed neuron/astrocyte cultures.

Briefly, day 12 NPCs were dissociated and replated to Matrigel-coated plates in STEMdiff astrocyte differentiation media (100-0013). Cultures were fed daily until day 18, then passaged and replated at 2.5 × 10^5^ cells/cm^2^. From day 18 to 35, cultures were fed every other day and passaged every 6 days at 2.5 × 105 cells/cm2. From day 36 through day 68, cultures were passaged and replated in STEMdiff astrocyte maturation media (100-0016). Cultures were fed every other day and passaged every 6 days at 2.5 × 10^5^ cells/cm^2^.

### Aβ Preparation

Amyloid-beta (Aβ_1-42_) was reconstituted and aggregated according to the Fujifilm/Cellular Dynamics Labeling Amyloid Beta protocol using the amyloid-beta aggregation kit (rPeptide). Briefly, lyophilized Aβ was reconstituted in 250 μL 10 mM NaOH and transferred to a 1.5 mL microcentrifuge tube. 250 μL of HPLC water was added, followed by 56 μL of 10X TBS (pH 7.4) and mixed by gentle pipetting. The Aβ solution was allowed to aggregate overnight in a normoxic incubator. The next day, the Aβ solution was centrifuged at 16,000x*g* for 1 minute to collect the aggregates. The supernatant was removed, and aggregates were washed with 200 μL 0.1 M NaHCO_3_ and resuspended by pipetting. Aggregates were washed an additional 4 times in Hank’s Balanced Salt Solution (HBSS) and resuspended in 200 μL HBSS, sonicated for 10 minutes, and stored in individual aliquots at -20C.

### Co-culture experiments

Day 38 iMGL were collected and replated onto day 68 astrocyte/neuron cultures in STEMdiff astrocyte maturation media supplemented with M-CSF (25 ng/ml), IL-34 (100 ng/ml),TGFβ-1 (50 ng/ml), CD200 (100 ng/mL) and CX3CL1 (100 ng/mL) at a ratio of 1:4. Cells were co-cultured for 72 hours prior to treatment with 50ug/ml oligomeric-Aβ or vehicle. After 24 hours, both adherent and suspension cell fractions were collected, washed with PBS, and dissociated with accutase. Cells were washed in wash buffer [PBS, 2% BSA, 0.2U/μl RNAse inhibitor] and centrifuged at 300xg for 5 min at 4°C, then filtered through 70 Flowmi strainers. Cells were counted using trypan blue on a Countess II (Invitrogen).

### Single-cell RNA sequencing and data processing

We prepared separate libraries from both the adherent and suspension cell fractions using 10x Genomics 3’ single-cell gene expression assays v3 and sequenced them on a NovaSeq 6000 to a depth of 50k reads/cell. Following sequencing and FASTQ generation, raw count matrices were produced using CellRanger v6.0.1. The resulting count matrices were corrected for ambient RNA using CellBender^66^ v0.3.0., and then processed using the Seurat v4.2.0 package in R. Cells containing < 2000 or > 9000 genes and/or > 7.5% mitochondrial RNA reads were removed. Data was scaled and log normalized, and principal component analysis (PCA) was performed on the top 2000 variable genes. We then used Seurat to perform UMAP dimension reduction on the top 15 PCs and used Seurat::findNeighbors and Seurat::findClusters to identify clusters based on a shared nearest neighbor (SNN) clustering algorithm with neighbors = 20 and Louvain resolution = 0.3. The data was manually inspected for cell doublets, and those containing mixed cluster signatures were removed, leaving 13,740 cells. The data processing steps and clustering procedure were repeated on the cleaned dataset. To identify marker genes for each cluster, we performed a Wilcoxon Rank Sum test with Bonferroni correction using Seurat::FindMarkers, testing each population against all other populations. Clusters were then annotated using canonical markers. To find genes differentially expressed between Aβ-treated and control cells, we used Seurat::FindMarkers to perform a Wilcoxon Rank Sum test with Bonferroni correction to compare Aβ treated cells against control cells within the clusters of interest. To further investigate iMGL, we subsetted the Aβ-treated iMGL and repeated the data scaling and normalization procedure using only those 3875 cells. We used Seurat to perform UMAP dimension reduction on the top 15 PCs and used Seurat::findNeighbors and Seurat::findClusters to identify clusters based on a shared nearest neighbor (SNN) clustering algorithm with neighbors = 20 and Louvain resolution = 0.3. To identify marker genes for each cluster, we performed a Wilcoxon Rank Sum test with Bonferroni correction using Seurat::FindMarkers, testing each iMGL cluster against all other iMGL populations. These cluster marker genes were used as genesets for GSEA.

## Supplementary Figures

**Supplementary Figure 1.**
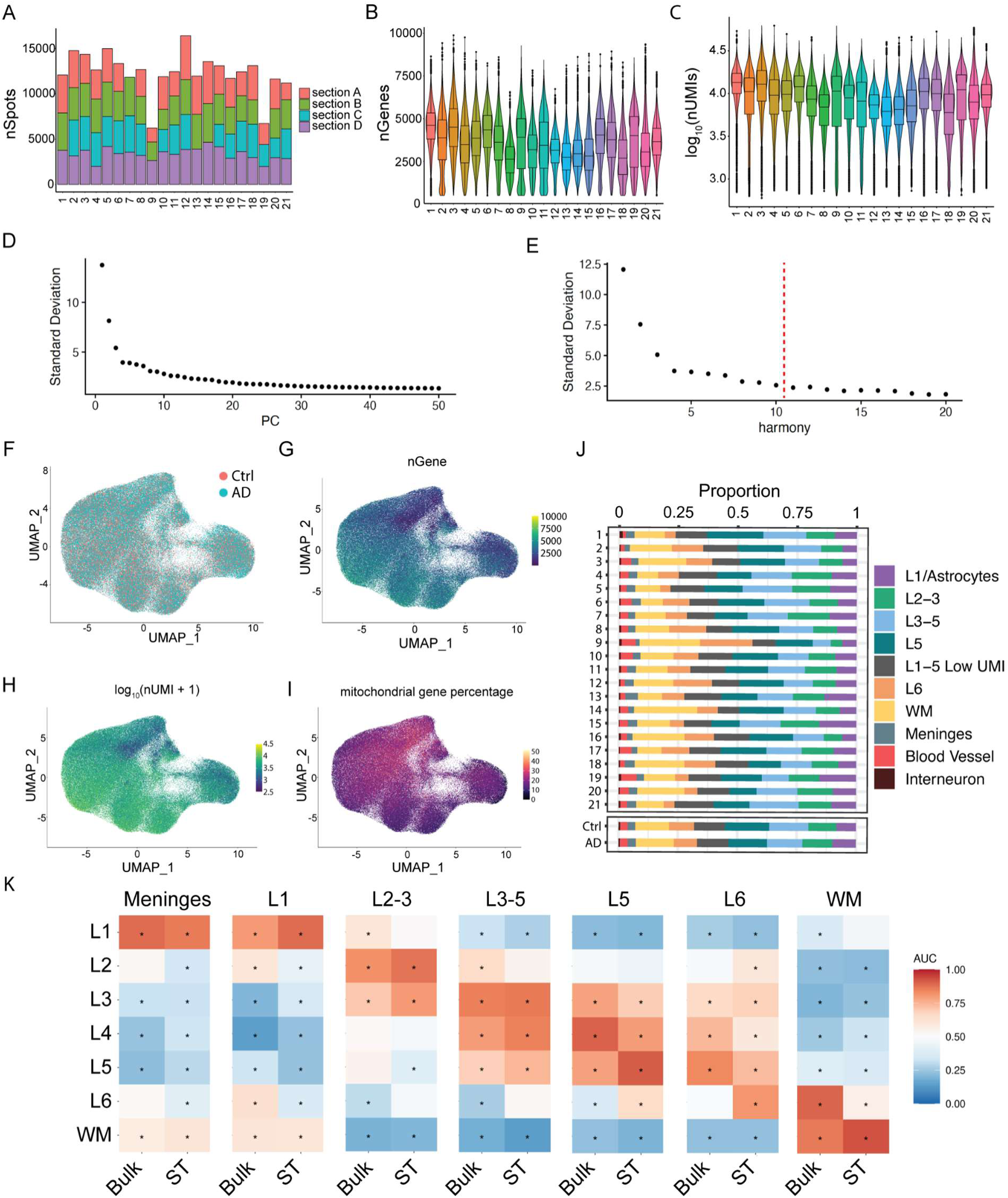
Spatial Transcriptomics QC. (A) Number of spots/sections analyzed for each individual. (B) Distribution of the number of detected genes per spot for each individual. (C) Distribution of the log-transformed number of detected UMIs for each individual. (D-E) Elbow plots showing the standard deviation for each principal component (D) or harmony embedding (E). (F-I) UMAP plots with spots colored by AD diagnosis (F), number of detected genes (G), number of detected UMIs (H), or mitochondrial gene percentage (I). (J) Proportion of spots represented by each spot cluster, by individual (top), or AD diagnosis (bottom). (K) Heatmap showing AUC values among cluster-enriched genes, comparing our cluster-enriched genes (columns) to cortical layer-enriched genes identified from previous studies^16–18^.

**Supplementary Figure 2.**
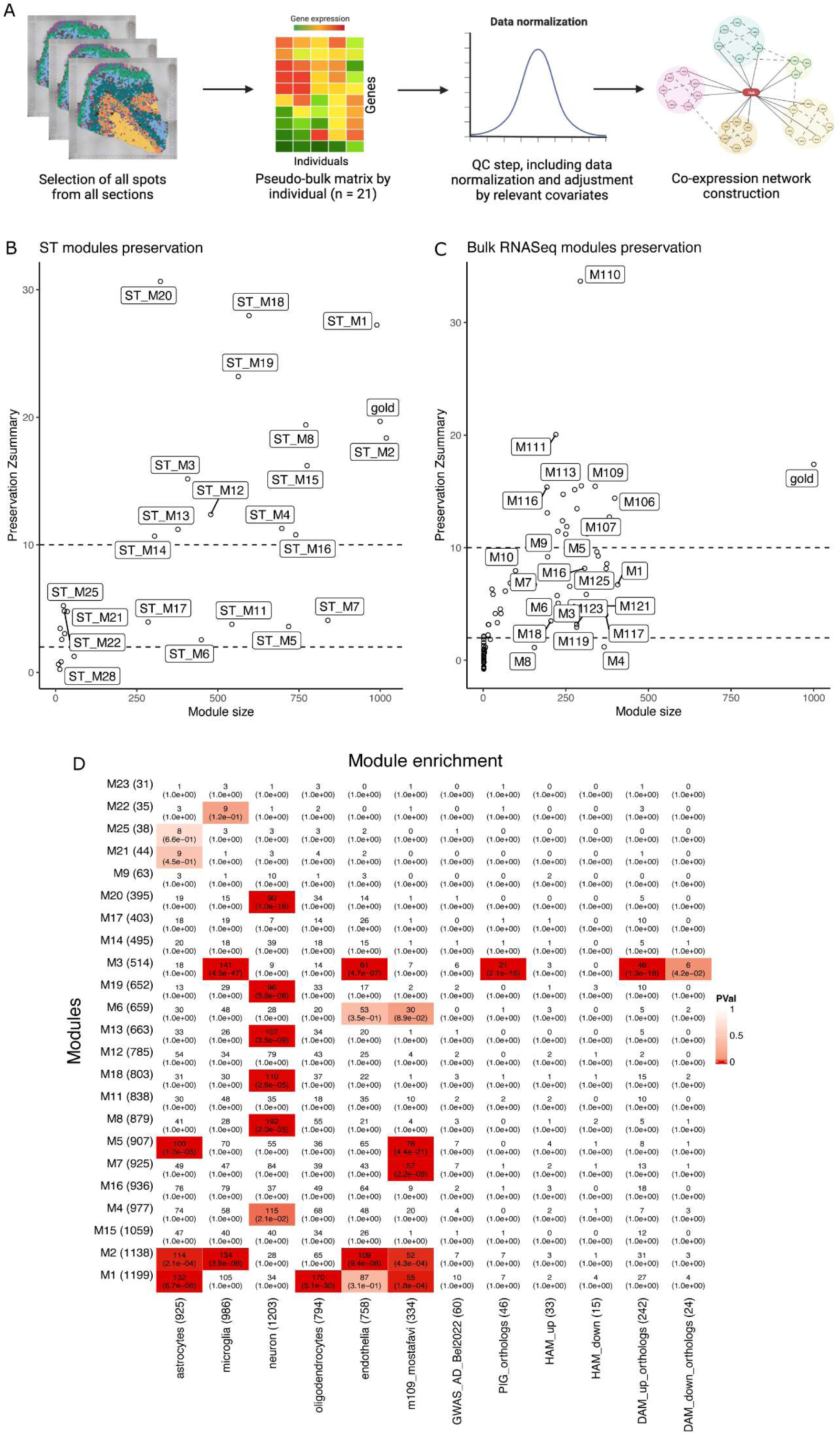
Spatial transcriptome network analysis. A) Spots were selected by the donor to generate a pseudo-bulk expression matrix. Following the quality control (QC) steps, the data was normalized and adjusted to remove technical confounders, and the final matrix was used as input to build the co-expression network. B-C) Composite module preservation statistics of the ST networks and previously published bulk RNASeq from the DLPFC region^11^. Plots show the summary statistics Z (y-axis) as a function of the module size (x-axis). Each dot represents a module labeled accordingly. The dashed lines indicate the Z = 2 and Z = 10 thresholds, respectively. B) The ST modules were used as reference. C) Bulk RNASeq DLPFC modules used as reference. D) Enrichment results between each module (y-axis) and the following gene lists on the x-axis: cell type marker genes for module 109^11^, AD GWAS genes^67^; plaque-induced genes (PIG orthologs)^26^; human-associated microglia (HAM)^7^; and damage-associated microglia (DAM)^27^. The number inside each cell represents gene overlap. P-values are from one-sided Fisher’s exact test followed by Bonferroni adjustment. The red color highlights the significant results (Adj p < 0.05).

**Supplementary Figure 3.**
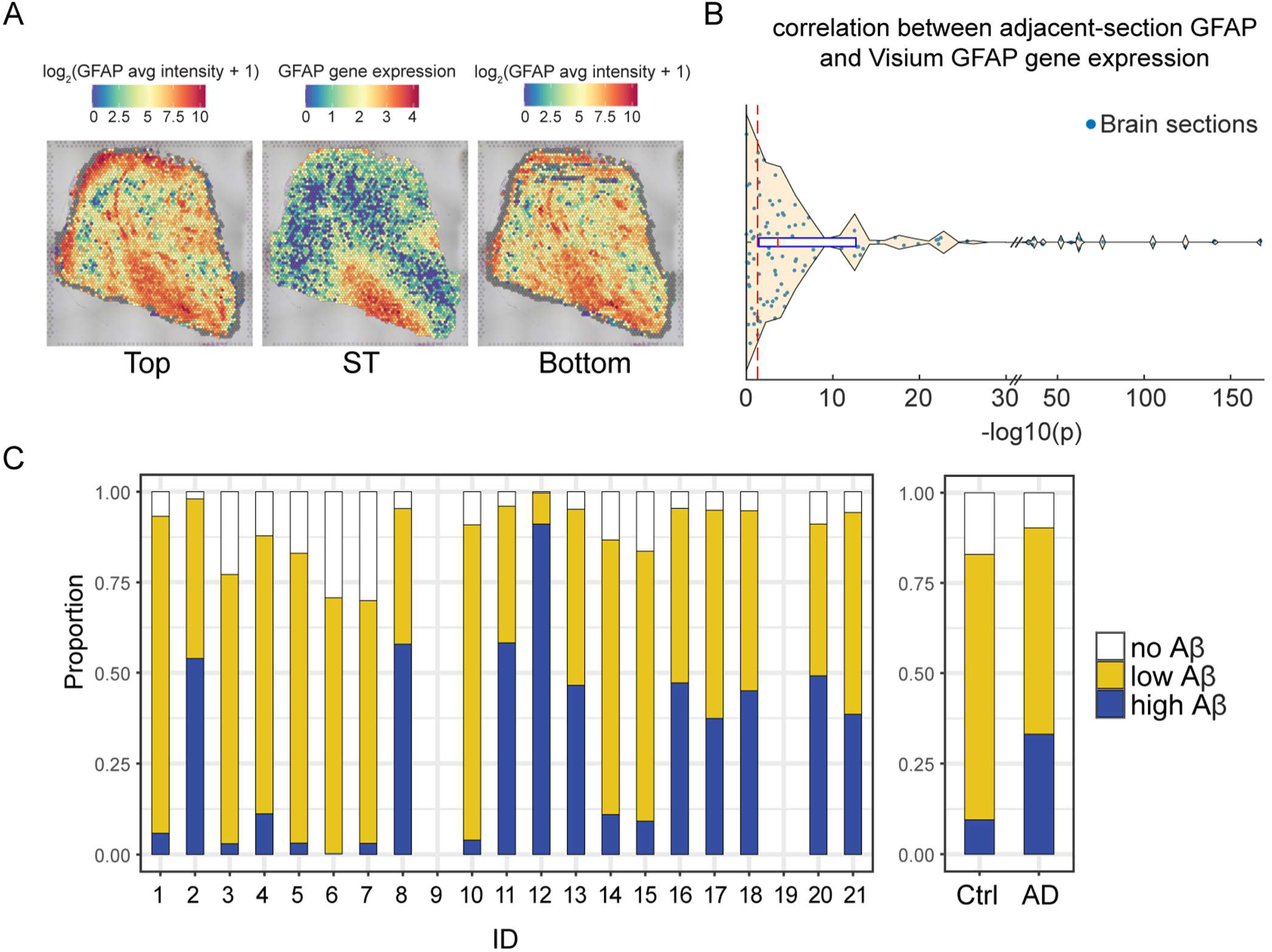
Validation of ST-adjacent IHC staining. (A) Feature plot showing per-spot GFAP gene expression (center) and average GFAP fluorescence intensities of adjacent sections (left, right) for a representative section (section 8_C). (B) For each ST-adjacent section, we correlated GFAP fluorescence intensity and GFAP gene expression across all tissue-covered spots. The distribution of -log10(p) across ST-adjacent sections displayed. (C) Stacked barplot showing the proportion of no, low, or high Aβ spots by individual (left), or clinical AD diagnosis (right).

**Supplementary Figure 4.**
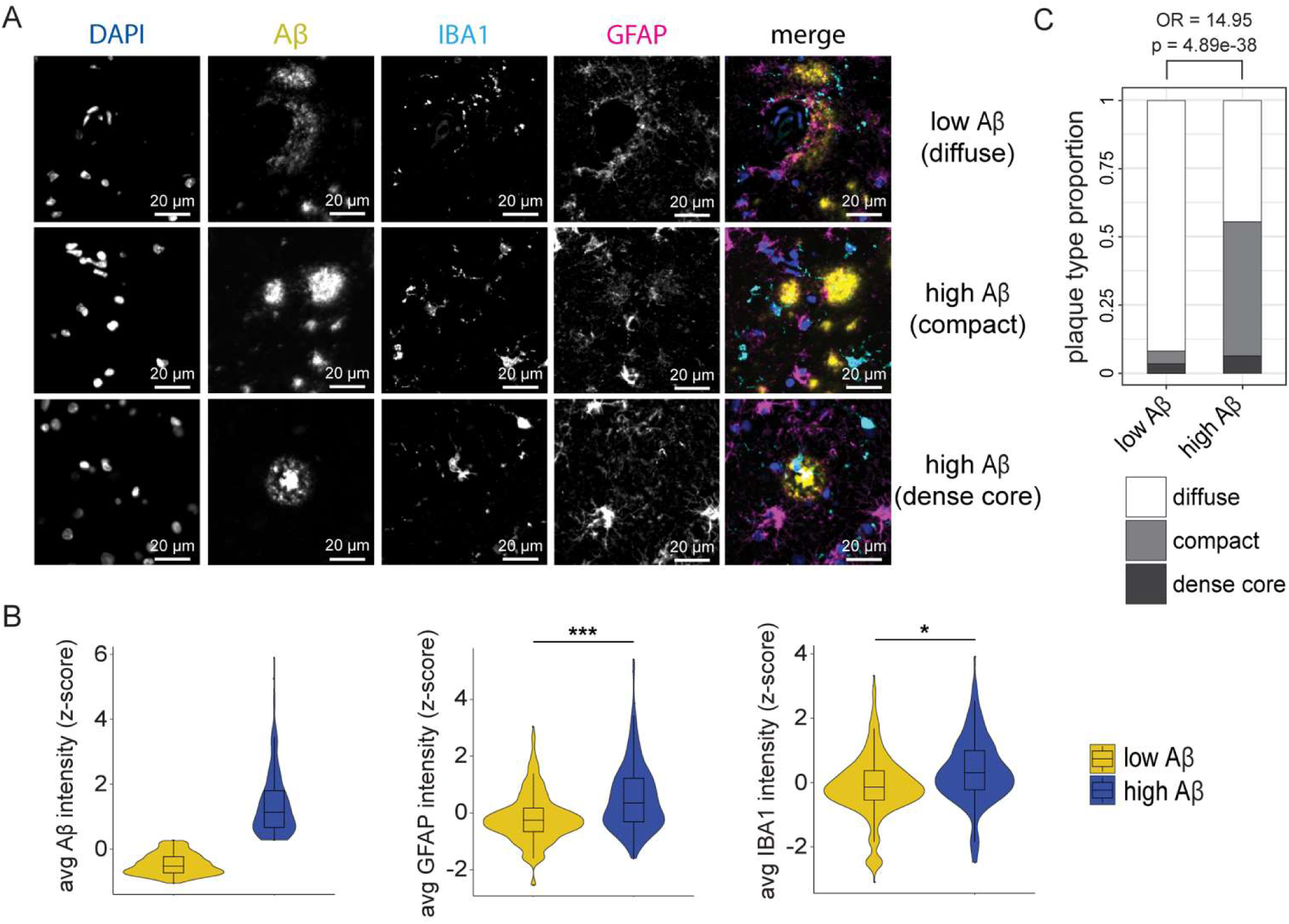
IHC of FFPE tissue. (A) Images of stained FFPE tissue sections from an individual with amyloid pathology, including a representative image of each major type of amyloid plaque (blue: DAPI, yellow: Aβ (4G8), cyan: IBA1, magenta: GFAP). (B) We manually annotated 722 plaques from 9 individuals, stratified them by low (bottom 75%) or high (top 25%) intensity of Aβ, then plotted the intensity distributions of Aβ (left), GFAP (middle), and IBA1 (right) (*p = 2.23e-08; ***p = 5.66e-18). (C) The proportion of plaque types from manually annotated plaques, stratified by Aβ intensity (OR: odds ratio).

**Supplementary Figure 5.**
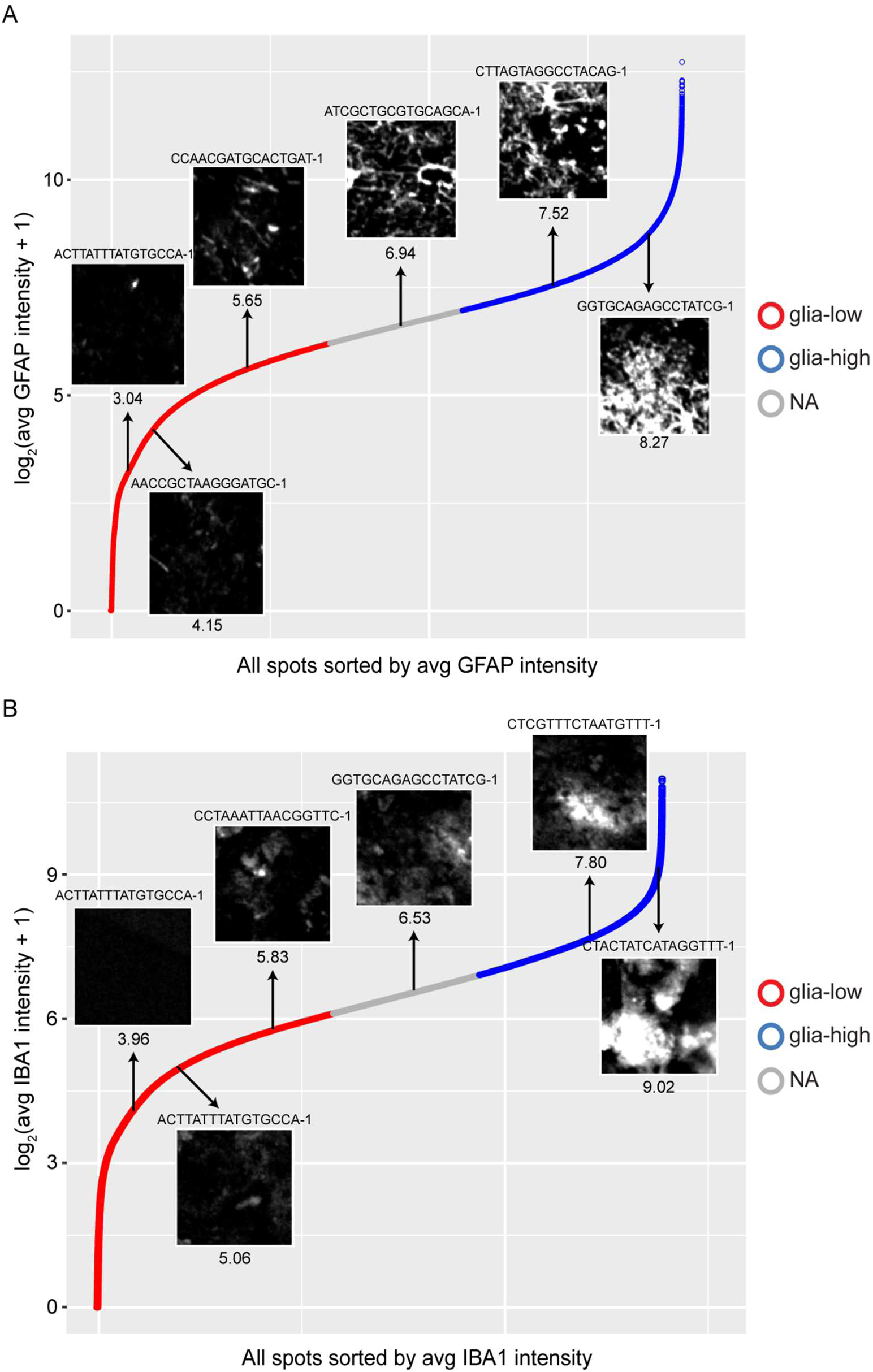
Representative images of GFAP and IBA1 from ST-adjacent spots. (A) All spots are plotted ordered by average GFAP intensity, and colored by spot stratification group (red: glia-low, blue: glia-high, gray: not assigned (NA)). Six representative spot images are shown from individual 11, labeled with their ST spot barcode (top) and log_2_(average GFAP intensity) (bottom). (B) All spots are plotted ordered by average IBA1 intensity, and colored by spot stratification group as in (A). Six representative spot images are shown from individual 11, labeled with their ST spot barcode (top) and log_2_(average IBA1 intensity) (bottom).

**Supplementary Figure 6.**
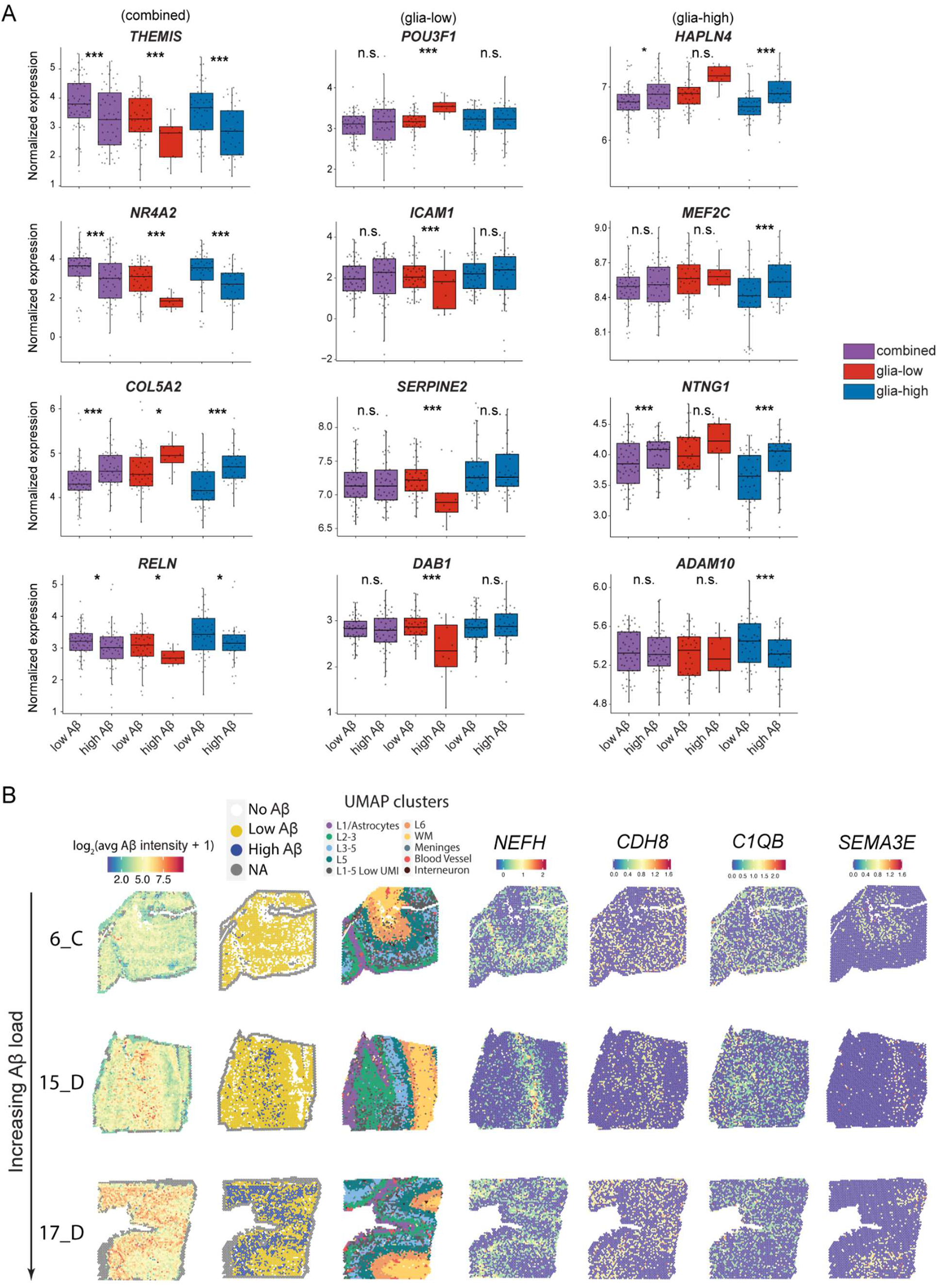
Additional characterization of the effects of low vs. high Aβ on transcriptome and apoptosis. (A) Boxplots of representative genes for each Aβ contrast (*p<0.05, ***p<5e-4, n.s.: not significant). (B) Columns from left to right correspond to spatial patterns of IF Aβ intensity, Aβ stratification group, UMAP labels, and expression levels of example genes.

**Supplementary Figure 7.**
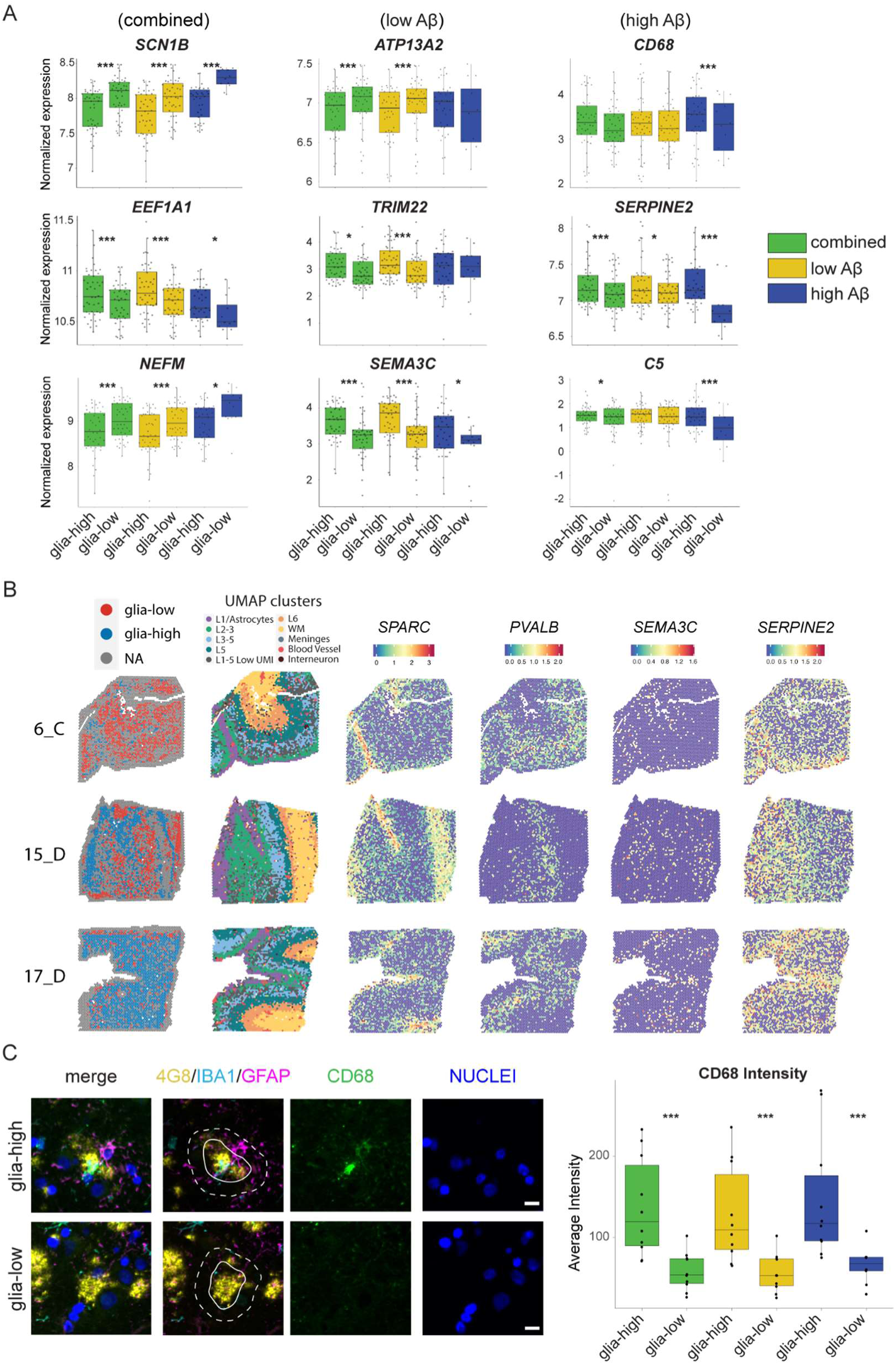
Additional characterization of the effects of glia on transcriptome. (A) Boxplots of representative genes for each Aβ contrast (*p<0.05, ***p<5e-4). (B) Columns from left to right correspond to spatial patterns of glia-high and glia-low microdomains, spot clusters, and expression levels of example genes. (C) Images of an FFPE tissue section stained with DAPI (blue), Aβ (4G8; yellow), IBA1 (cyan), GFAP (magenta), and CD68 (green) (Scale bar = 25 µm). CD68 average intensity was quantified in the area surrounding plaques (25 µm ring, between the solid and dotted lines in representative images; *** = p<1e-20).

**Supplementary Figure 8.**
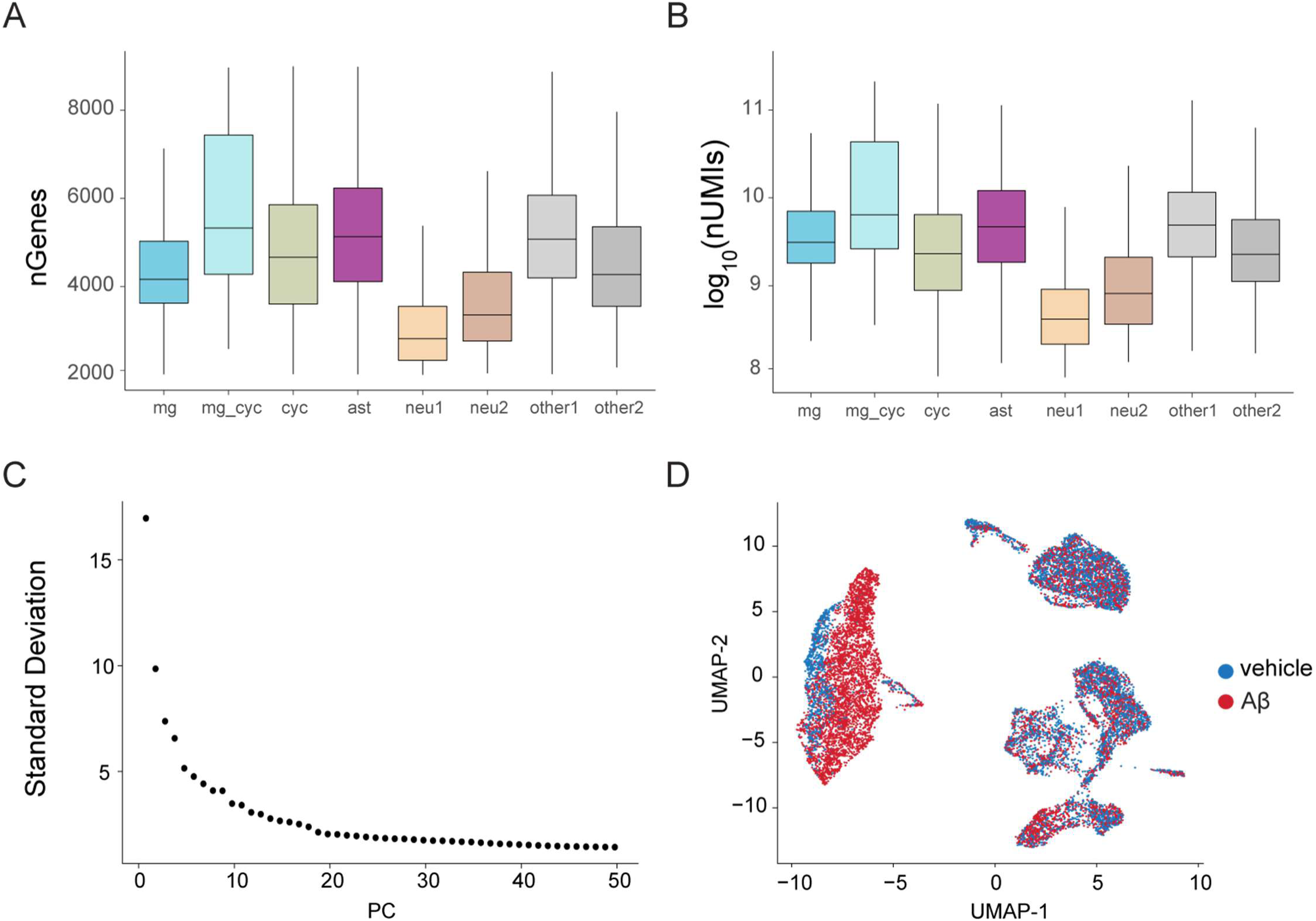
Single-cell RNA-seq QC plots. (A and B) Distributions of numbers of detected genes (A) and log_10_(nUMIs) (B) for each major cell type. (C) Elbow plot showing the standard deviation attributed to each PC. The top 15 PCs were used for UMAP dimension reduction. (D) UMAP plot with cells colored by treatment with vehicle (blue) or Aβ oligomers (red).

## Supplementary Tables

Supplementary Table 1. Metadata for 21 ROSMAP participants analyzed by ST

Supplementary Table 2. Enriched genes for each ST spot cluster

Supplementary Table 3. GSEA of cell2location cell abundance estimates in each brain region

Supplementary Table 4. Genes and pathways associated with each ST gene module

Supplementary Table 5. Manual plaque type annotation for a subset of ST spots

Supplementary Table 6. Differential genes and pathways for low vs. high Aβ contrasts

Supplementary Table 7. Cell2location cell abundance estimates for each contrast between stratified spot groups

Supplementary Table 8. Ligand-Receptor differential expression (NICHES) for each contrast between stratified spot groups

Supplementary Table 9. Differential genes and pathways for glia-high vs. glia-low contrasts

Supplementary Table 10. Differential genes and pathways for Aβ-responses in *in vitro* co-cultured glia

Supplementary Table 11. Differential genes for iMGL Aβ clusters

Supplementary Table 12. Differential genes for iMGL Aβ clusters contrasted to control iMGL

Supplementary Table 13. Key Resources Table

## Notes

### Competing Interest Statement

The authors have declared no competing interest.

